# *Chlamydia muridarum* Causes Persistent Subclinical Infection and Elicits Innate and Adaptive Immune Responses in C57BL/6J, BALB/cJ and J:ARC(S) Mice Following Exposure to Shedding Mice

**DOI:** 10.1101/2024.07.16.603732

**Authors:** Noah Mishkin, Sebastian E Carrasco, Michael Palillo, Panagiota Momtsios, Cheryl Woods, Kenneth S Henderson, Ana Leda F Longhini, Chelsea Otis, Rui Gardner, Ann M Joseph, Gregory F Sonnenberg, Jack Palillo, Rodolfo J Ricart Arbona, Neil S Lipman

## Abstract

*Chlamydia muridarum* (Cm) has reemerged as a moderately prevalent infectious agent in research mouse colonies. Despite its’ experimental use, few studies evaluate Cm’s effects on immunocompetent mice following its natural route of infection. A Cm field isolate was administered (orogastric gavage) to 8-week-old female BALB/cJ (C) mice. After confirming shedding (through 95d), these mice were cohoused with naïve C57BL/6J (B6), C, and Swiss (J:ARC[S]) mice (n=28/strain) for 30 days. Cohoused mice (n=3-6 exposed and 1-6 control/strain) were evaluated 7, 14, 21, 63, 120, and 180 days post-cohousing (DPC) via hemograms, serum biochemistry analysis, fecal qPCR, histopathology, and Cm MOMP immunohistochemistry. Immunophenotyping was performed on spleen (B6, C, S; n=6/strain) and intestines (B6; n=6) at 14 and 63 DPC. Serum cytokine concentrations were measured (B6; n=6 exposed and 2 control) at 14 and 63 DPC. All B6 mice were shedding Cm by 3 through 180 DPI. One of 3 C and 1 of 6 S mice began shedding Cm at 3 and 14 DPC, respectively, with the remaining shedding thereafter. Clinical pathology was nonremarkable. Minimal-to-moderate enterotyphlocolitis and gastrointestinal associated lymphoid tissue (GALT) hyperplasia was observed in 15 and 47 of 76 Cm-infected mice, respectively. Cm antigen was frequently detected in GALT-associated surface intestinal epithelial cells. Splenic immunophenotyping revealed increased monocytes and shifts in T cell population subsets in all strains/timepoints. Gastrointestinal immunophenotyping (B6) revealed sustained increases in total inflammatory cells and elevated cytokine production in innate lymphoid cells and effector T cells (large intestine). Elevated concentrations of pro-inflammatory cytokines were detected in the serum (B6). Results demonstrate that while clinical disease was not appreciated, 3 commonly utilized strains of mice are susceptible to chronic enteric Cm infection which may alter various immune responses. Considering the widespread use of mice to model GI disease, institutions should consider excluding Cm from their colonies.

## Introduction

The murine bacterial pathogen *Chlamydia muridarum* (Cm) has recently reemerged as an opportunist affecting research mouse colonies across the United States and Europe.^35,48,52^ The re-emergence of Cm is concerning as, based on historical reports and experimental use, Cm infection has the potential to cause clinical disease, elicit immune responses, and may otherwise confound research data.^4,10,34,35,51^ Initially identified in the 1930’s and 1940’s in laboratory mice used to study human viral respiratory disease, Cm was recognized as an intracellular agent capable of causing pulmonary disease.^9,18,37^ Subsequently, Cm has been utilized extensively to model human *Chlamydia trachomatis* infection in mice.^10,42^ Despite its long-standing historical significance and experimental use, there is a limited understanding of the impact of naturally acquired Cm infection on immunocompetent mouse strains.^34^

Cm, the only Chlamydial species infecting mice, is an obligate intracellular gram-negative bacterium with tropism for mucosal epithelium.^42^ Both the nonreplicating and infectious elementary bodies (EB) and the replicating and noninfectious reticulate bodies (RB) may be visualized microscopically within intracellular inclusion bodies (IB).^42^ Cm infection in immunocompetent mouse strains results in transient local pathology or asymptomatic shedding, while immunodeficient strains develop and often succumb to fulminant disease.^25,30,43,66^ Immunocompetent mice were thought to resolve infection, however recent experimental studies have demonstrated that Cm can cause chronic, subclinical infections in which mice remain culture positive and/or IBs are observed microscopically for extended periods (exceeding 100 days post-infection in one study, and 260 days in another).^22,66^ Differences in response to experimental Cm infection have been reported in immunocompetent strains, including but not limited to C57BL/6 (B6), BALB/c (C), and C3H mice.^42^ Of these, B6 mice generally sustain lower Cm burdens, 50-100% shorter durations of infection, and lesser respiratory pathology as compared to C mice, indicating a T helper (Th) 1 cell immune response can better control Cm infection.^25, 42, 66^ Experimental infection of immunodeficient strains, including athymic nude, SCID, and other immunodeficient transgenic strains, such as interferon-γ (IFNγ), toll-like receptor (TLR), STAT1, MHCII/CD4, and Rag1 knockouts, have been shown to cause severe clinical signs, widespread tissue distribution, and a protracted course. The clinical signs, tissue distribution, and kinetics of Cm infection vary based on mouse strain, infectious dose of Cm, and the route of inoculation. Commonly observed phenotypes include include significant inflammation, fatal pulmonary disease, and dissemination to sites other than the site of administration.^5,6,13,30,41,42^

While experimental use of Cm demonstrated the potential for chronic infection of immunocompetent strains, these studies generally utilized atypical routes of inoculation (urogenital and/or upper respiratory tract), high doses of inoculum (10^5^-10^7^ inclusion forming units [IFUs]), as well as pretreatment with progesterone.^42^ Recently, a study investigating the ability of Cm to persist in vivo demonstrated long-term colonization of the gastrointestinal tract at a lower oral dose of 10^3^ IFU, indicating an ID_50_ at least 100-fold lower than had been used historically.^66^ Furthermore, experimental infections in immunocompetent and immunodeficient mouse strains challenged with Cm Nigg strains via orogastric gavage have shown that this Cm persisted in the cecum and colon and used chromosome-encoded proteins to evade interferon-γ-producing type 3 innate lymphoid cells to establish long-lasting colonization of the gastrointestinal tract.

The recent identification of Cm in 2 genetically engineered mouse (GEM) colonies following an investigation into the etiology of unexpected pulmonary inflammatory lesions led to the determination of its’ considerable intra- and inter-institutional prevalence in research mouse colonies. This investigation revealed that 63% of institutional soiled bedding sentinels (97 rooms surveyed), 33% of 58 incoming mouse shipments from 39 academic institutions, 3% of 11,387 microbiota samples submitted from 120 institutions (with 14% of institutions yielding at least a single positive result), and 16% of 900 diagnostic samples submitted from 96 institutions were Cm positive.^35^ This surprising prevalence is concerning as subclinical infection with the bacterium has the ability to confound studies in which infected mice are utilized, and Cm can cause clinical disease in NOD.Cg-*Prkdc*^scid^ Il2rg^tm1Wjl^/SzJ (NSG) mice and GEM strains following unwitting exposure through the bacterium’s presumptive natural route of infection.^34,35,51^ NSG mice used as contact sentinels developed severe clinical signs including lethargy, dyspnea, unkempt coats, and weight loss resulting from a moderate-to-severe bronchointerstitial pneumonia and extensive colonization of the small and large intestines.^51^ Additionally, 2 GEM strains with impaired IFNγ signaling and Th1 cell responses presented with clinical disease following spontaneous Cm infection.^34^

Despite Cm’s widespread use in the laboratory, there is a paucity of information pertaining to the biology of Cm in immunocompetent laboratory mice following exposure by its’ presumptive natural oral route and dose of infection. Given the potential for Cm infection to alter immunological and other biological processes in experimental models, its’ prevalence in research colonies, and clinical disease in immunocompromised and GEM strains, there is a need to better understand the biology of Cm in natural/spontaneous infections. The studies described herein were conducted to characterize the clinical disease and pathology, tissue tropism, fecal shedding, and immune responses following exposure of 3 immunocompetent strains/stocks to Cm-shedding mice. The principal study aim was to better understand Cm’s potential impact on studies, particularly those investigating or dependent on gastrointestinal tract physiology or immunology. This understanding is crucial to determine whether Cm should be excluded from research colonies, how to best identify infected animals, and provide foundational data necessary to design future studies on how best to eradicate Cm from infected colonies.

## Materials and Methods

### Investigative plan (Figure 1)

To explore clinical disease, course of infection, tissue tropism, pathology, fecal shedding kinetics, and the immunologic effects of Cm acquired through natural routes of transmission, we first established a cohort of chronically Cm-infected female C mice. C mice were utilized given experimental data describing their greater Cm burden and prolonged fecal shedding.^66^ After confirming these mice were chronically shedding Cm through day 95 (via fecal qPCR), the mice were then cohoused with naïve BALB/c (C), C57BL6 (B6) and J:ARC(S) [S] mice (1 infected C mouse/4 uninfected C, B6, or S mice/cage; n=28/strain) to establish infection through the fecal-oral and/or the inhalational routes. Inoculated C mice were then removed after 30 days. C and B6 mice were selected given their differing Th1 and Th2 immune responses, colonization differences observed in experimental studies, and their common use as research models, both as inbred as well as the background strain of many genetically engineered models. The outbred S mouse was evaluated to examine Cm’s pathobiology in a mouse with greater genetic diversity. Control C, B6, and S mice (n=16/strain) were cohoused with a naïve, Cm-free C mouse as was described for the experimental mice.

**Figure 1.**
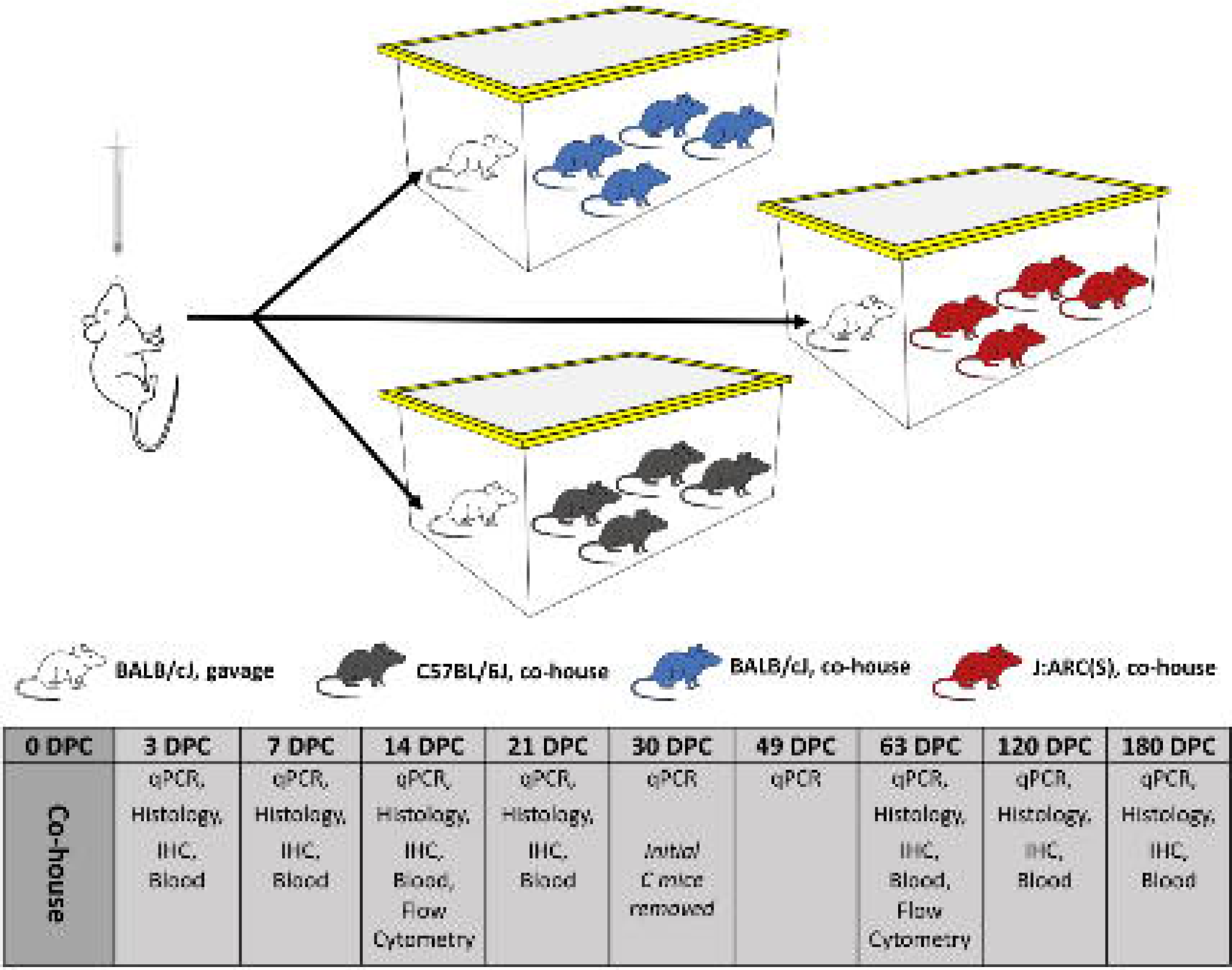
Investigative plan. BALB/cJ (C) mice inoculated with and shedding Cm were cohoused with naïve C, C57BL/6J (B6), or J:ARC(S) [S] mice (1 infected C mouse with 4 naïve C, B6, or S mice per cage) for 30 days. Mice were monitored for 180 days. Three experimental and 1 control mouse per strain/stock were selected for qPCR, hemogram/serum biochemistry analysis, complete necropsy and IHC at 3, 7, 14, 21, 63, 120, and 180 DPC. Additional mice were sacrificed at 14 and 63 DPC for splenic and gastrointestinal tract (B6 only) immunophenotyping.

Following cohousing, the C, B6, and S experimental and control mice were monitored daily for activity, respiratory rate and effort, coat condition and posture, and body condition score (BCS) for up to 180 days after placement of the Cm-shedding or Cm-free C cohoused mouse. On 3, 7, 14, 21, 30, 49 (feces only), 63, 120, and 180 days post-cohousing (DPC), 3 experimental and 1 control mouse per strain were euthanized and blood was collected for hemogram and serum biochemistry analysis, feces were collected for qPCR, and a complete necropsy was performed; additional mice (n=6 experimental and control mice/strain) were assessed at 14 and 63 DPC. Immunohistochemistry (IHC) for chlamydial major outer membrane protein (MOMP) was performed on all tissues assessed, and in-situ hybridization (ISH) for Cm nucleic acid was performed on select tissues (lungs and gastrointestinal tract) from a subset of mice (n=2 B6 and n=7 S mice) when MOMP signal was absent or inconclusive in Cm qPCR+ mice. To assess the systemic and local immunologic response to Cm infection, immunophenotyping by flow cytometry of the spleen (n=6 infected and control mice/strain/timepoint; C, B6 and S) and gastrointestinal tract (n=6 infected and control mice/timepoint; B6 only) specific immune responses were characterized at early (14 DPC) and late (63 DPC) timepoints. Select serum cytokine levels were also assessed 14 and 63 DPC (n=5 infected and n=2 control/timepoint) in B6 mice.

A subset of male C, B6 and S mice (n=5/strain) were also inoculated with Cm via orogastric gavage to investigate potential sex differences and the colonization of the male reproductive tract.

### Animals

Six- to 8-week-old C57BL/6J (B6; n=58F, 5M), BALB/cJ (C; n=81F, 5M), and J:ARC(S) (S; n=46F, 5M; The Jackson Laboratory, Bar Harbor, ME) mice were used. All mice were received free of ectromelia virus, Theiler meningoencephalitis virus (TMEV), Hantaan virus, K virus, LDH elevating virus (LDEV), lymphocytic choriomeningitis virus (LCMV), mouse adenovirus (MAV), murine cytomegalovirus (MCMV), murine chapparvovirus (MuCPV), mouse hepatitis virus (MHV), minute virus of mice (MVM), murine norovirus (MNV), mouse parvovirus (MPV), mouse thymic virus (MTV), pneumonia virus of mice (PVM), polyoma virus, reovirus type 3, epizootic diarrhea of infant mice (mouse rotavirus, EDIM), Sendai virus, murine astrovirus-2 (MuAstV-2), *Bordetella* spp., *Citrobacter rodentium*, *Clostridium piliforme*, *Corynebacterium bovis*, *Corynebacterium kutscheri, Filobacterium rodentium* (CAR bacillus), *Mycoplasma pulmonis* and other *Mycoplasma* spp., *Salmonella* spp., *Streptobacillus moniliformis, Helicobacter* spp., *Klebsiella pneumoniae* and *K. oxytoca*, *Pasteurella multocida*, *Rodentibacter pneumotropicus/heylii*, *Pseudomonas aeruginosa*, *Staphylococcus aureus*, *Streptococcus pneumoniae* and Beta-hemolytic *Streptococcus* spp., *Yersinia enterocolitica* and *Y. pseudotuberculosis*, *Proteus mirabilis*, *Pneumocystis murina*, *Encephalitozoon cuniculi*, ectoparasites (fleas, lice, and mites), endoparasites (tapeworms, pinworms, and other helminths), protozoa (including *Giardia* spp. and *Spironucleus* spp.), *Toxoplasma gondii*, trichomonads, and dematophytes. Additionally, all mice were confirmed *Chlamydia muridarum* negative prior to initiating studies.

### Husbandry and housing

Mice were maintained in autoclaved, individually ventilated, polysulfone shoebox cages with stainless-steel wire-bar lids and filter tops (IVC; no. 19, Thoren Caging Systems, Inc., Hazelton, PA) on autoclaved aspen chip bedding (PWI Industries, Quebec, Canada) at a density of no greater than 5 mice per cage. Each cage was provided with a Glatfelter paper bag containing 6 g of crinkled paper strips (EnviroPak, WF Fisher and Son, Branchburg, NJ) and a 2-inch square of pulped virgin cotton fiber (Nestlet, Ancare, Bellmore, NY) for enrichment. Mice were fed a natural ingredient, closed source, γ-irradiated, autoclaved feed (LabDiet 5KA1, PMI, St. Louis, MO) ad libitum^57^ All animals were provided autoclaved reverse osmosis acidified (pH 2.5 to 2.8 with hydrochloric acid) water in polyphenylsulfone bottles with stainless-steel caps and sipper tubes (Techniplast, West Chester, PA). Cages were changed every 7 days within a class II, type A2 biological safety cabinet (LabGard S602-500, Nuaire, Plymouth, MN). All cages were housed within a dedicated, restricted-access cubicle, which was maintained on a 12:12-h light:dark cycle (on 6 AM; off 6 PM), relative humidity of 30 to 70%, and room temperature of 72 ± 2°F (22.2 ± 1.1°C). All cage changing and animal handling was performed by the first author (NM). The animal care and use program at Memorial Sloan Kettering Cancer Center (MSK) is accredited by AAALAC International and all animals are maintained in accordance with the recommendations provided in the Guide.^23^ All animal use described in this investigation was approved by MSK’s IACUC in agreement with AALAS’ position statements on the Humane Care and Use of Laboratory Animals and Alleviating Pain and Distress in Laboratory Animals.^1,2^

### Generation of Cm-shedding C mice

Cm qPCR negative C mice were individually identified, randomized (randomizer.org) and assigned to either the uninfected control (n=10) or infected (n=25) groups. For the latter, each mouse was inoculated with 2.72 x 10^3^ IFU of a previously isolated Cm field strain in 100 µL sucrose-phosphate-glutamic acid buffer (SPG, pH 7.2) via orogastric gavage.^35^ A new, sterilized gavage needle (22g x 38.1mm, Cadence Science, Cranston, RI) was used for each cage. Control mice were maintained in an IVC with no intervention other than fecal sampling.

An infected mouse was euthanized via CO_2_ asphyxiation on days 7, 14, and 21 days post-inoculation DPI to confirm infection by qPCR, and histology and immunohistochemistry were conducted to determine on which tissues to focus subsequent histologic and immunohistochemical analyses.

Fecal pellets (10-12 pellets pooled by cage/sample; n = 5 mice/cage) were collected on 7, 30, 75, and 95 days post-infection (DPI) and evaluated by qPCR to confirm initial and sustained Cm infection and lack thereof (controls). On confirmation of sustained infection at 95 DPI, Cm qPCR negative B6, C, and S mice were similarly identified, randomized, and assigned to either the uninfected control (n=18/strain) or infected (n=28/strain) groups, and cohoused with either Cm-shedding or Cm-free C mice as described in the investigative plan.

### Clinical monitoring

All mice were monitored daily to assess activity, respiratory rate and effort, coat condition and posture, and BCS.^58^ Animals displaying weight loss (20% reduction from baseline) or reduced body condition (BCS < 2/5), presenting with dyspnea and/or cyanosis, and/or failing to respond to stimulation were considered to have met the study’s humane endpoint necessitating euthanasia.

### Fecal collection

Fecal pellets for qPCR were collected ante- and post-mortem. When collected antemortem, the mouse was lifted by the base of the tail and allowed to grasp onto a wire-bar lid while a sterile 1.5 mL microcentrifuge tube was placed underneath the anus for fecal collection directly into tube. If at the end of a 30 second period the animal did not defecate, it was returned to the cage and allowed to rest for at least 2 minutes and collection reattempted until a sample was produced. Postmortem fecal collection was collected with aseptic technique directly from the colon. Individual fecal pellets were either pooled for submission (non-terminal collection for inoculated mice at all timepoints and for cohoused mice at 30 and 49 DPC), or submitted directly (for all euthanized and necropsied mice.)

### Chlamydia muridarum qPCR assay and amplicon sequencing

Total nucleic acid from feces from Cm-infected and control mice were extracted using a magnetic-based total nucleic acid isolation kit (MagMAX Total Nucleic Acid Isolation Kit, Life Technologies, Carlsbad, CA) following internally optimized protocols. Quantitative (real-time) Polymerase Chain Reaction (qPCR) amplification was performed using the ThermoFisher qPCR Systems (Waltham, MA) according to the manufacturer’s instructions. A proprietary real-time fluorogenic 5’ nuclease PCR assay specifically targeting a 425 bp fragment of Cm 23S rRNA was used to determine the presence of chlamydial DNA in samples as previously described.^35^ Samples that amplified during initial testing were subsequently retested to confirm the original finding. To monitor sample suitability (e.g., evaluating successful DNA recovery after extraction and assessing whether PCR inhibitors were present), an exogenous nucleic acid recovery control assay was added to each sample prior to magnetic nucleic acid isolation. A second real-time fluorogenic 5′ nuclease PCR assay was used to target the exogenous template to serve as a sample suitability control and was performed simultaneously with the Cm assay. A 100-copy/reaction positive control plasmid template containing the Cm target template was co-PCR amplified with the test sample to demonstrate master mix and PCR amplification equipment function. The double-stranded nucleotide sequences of selected purified PCR products were determined using Sanger dye-termination sequencing (Genomics Core, Tufts University Medical School, Boston, MA).^20^ Cm copies per reaction in a sample were estimated by comparing the average sample and 100-copy positive template control cycle-threshold values; a difference of 3.3 Ct corresponds to approximately a 10-fold difference in copy number.^60^

### Clinical and anatomic pathology

Following euthanasia via CO2 asphyxiation, peripheral blood was collected by cardiac puncture and/or from the caudal vena cava; bilateral thoracotomy was performed immediately after. For hematology, blood was collected into tubes containing EDTA (BD Microtainer K2EDTA, Becton, Dickinson and Company, Franklin Lakes, NJ). Analysis was performed on an automated hematology analyzer (IDEXX Procyte DX, Columbia, MO) and the following parameters determined: white blood cell count, red blood cell count, hemoglobin concentration, hematocrit, mean corpuscular volume, mean corpuscular hemoglobin, mean corpuscular hemoglobin concentration, red blood cell distribution width standard deviation and coefficient of variance, reticulocyte relative and absolute counts, platelet count, platelet distribution width, mean platelet volume, and relative and absolute counts of neutrophils, lymphocytes, monocytes, eosinophils, and basophils. Manual differential counts were performed in samples with read errors. For serum biochemistry analysis, blood was collected into tubes containing a serum separator (BD Microtainer SST, Becton, Dickinson and Company), the tubes were centrifuged, and the serum analyzed. An automated analyzer (Beckman Coulter AU680, Brea, CA) was used to determine the concentration of the following analytes: alkaline phosphatase, alanine aminotransferase, aspartate aminotransferase, creatine kinase, gamma-glutamyl transpeptidase, albumin, total protein, globulin, total bilirubin, blood urea nitrogen, creatinine, cholesterol, triglycerides, glucose, calcium, phosphorus, chloride, potassium, and sodium. The Na/K and the albumin/globulin ratios were calculated. Complete blood count and serum biochemistry values were compared to in-house reference values from healthy, experimentally naïve, adult C, B6 and S mice.

A complete necropsy was then performed, and gross lesions recorded. Tissues including heart, thymus, lungs, liver, gallbladder, kidneys, pancreas, stomach, duodenum, jejunum, ileum, cecum, colon, lymph nodes (mandibular, mesenteric), salivary glands, skin (trunk and head), urinary bladder, uterus, cervix, vagina, ovaries, oviducts (or testes, epididymides, seminal vesicles, and prostate for male cohort), adrenal glands, spleen, thyroid gland, esophagus, trachea, spinal cord, vertebrae, sternum, femur, tibia, stifle joint, skeletal muscle, nerves, skull, nasal cavity, oral cavity, teeth, ears, eyes, pituitary gland, and brain were fixed in 10% neutral buffered formalin for at least 72h. The entire length of the intestines were processed into Swiss rolls. After fixation, the skull, spinal column, sternum, femur and tibia were decalcified in a 12-15% EDTA solution prepared in-house. Representative tissue sections were then processed in ethanol and xylene and embedded in paraffin in a tissue processor (Leica ASP6025, Leica Biosystems, Deer Park, IL). Paraffin blocks were sectioned at 5 μm, stained with hematoxylin and eosin (H&E), and examined by a board-certified veterinary pathologist (SC).

Gastrointestinal lymphoid tissue (GALT) hyperplasia was assessed in all mice evaluated histologically. Briefly, a scoring system was used: 0 - normal; 1 - mild evidence of follicular hyperplasia (e.g., focal-to-multifocal areas of increased lymphoid cellularity with no displacement or rare outpouching of the serosal surface); 2 - moderate evidence of follicular hyperplasia (e.g., multifocal-to-diffuse increased lymphoid cellularity with some displacement/outpouching of the serosal surface, occasional prominent germinal centers); and, 3 - marked evidence of follicular hyperplasia (e.g., diffusely increased lymphoid cellularity with frequent displacement/outpouching of the serosal surface, frequent/extensive prominent germinal centers). Examples of each score are provided in Figure 4A.

Immunohistochemistry (IHC) - All tissues collected from both mice inoculated via orogastric gavage (n=3 C mice) and mice exposed to Cm through cohousing at 3, 7, and 14 DPC (n=12 mice/strain) were processed and stained for *Chlamydia* MOMP using a technique optimized and validated by MSK’s Laboratory of Comparative Pathology.^16^ Thereafter, at 21, 63, 120, and 180 DPC (n=16 mice/strain), only the gastrointestinal, reproductive tract and pulmonary tissues collected from cohoused mice were assessed.

Briefly, formalin-fixed, paraffin-embedded sections were stained using an automated staining platform (Leica Bond RX, Leica Biosystems). Following deparaffinization and heat-induced epitope retrieval in a citrate buffer at pH 6.0, the primary antibody against *Chlamydia* MOMP (NB100-65054, Novus Biologicals, Centennial, CO) was applied at a dilution of 1:500. A rabbit anti-goat secondary antibody (Cat. No. BA-5000, Vector Laboratories, Burlingame, CA) and a polymer detection system (DS9800, Novocastra Bond Polymer Refine Detection, Leica Biosystems) was then applied to the tissues. The 3,3’-diaminobenzidine tetrachloride (DAB) was used as the chromogen, and the sections were counterstained with hematoxylin and examined by light microscopy. Reproductive tracts from TLR3- deficient mice experimentally infected with *Chlamydia muridarum* strain Nigg were used as a positive control.^51^ Positive MOMP immunolabeling was identified as discrete, punctate chromogenic brown dots under bright field microscopy.

All gastrointestinal tracts evaluated via IHC were assigned a score associating Cm MOMP antigen immunostaining of surface small intestinal, cecal, and colonic epithelial cells with GALT at 3, 7, 14, 21, 63, 120, and 180 DPC: 0 - no association (MOMP immunostaining was either not observed or was exclusively in epithelial cells without GALT association); 1 -occasional association (1-49% immunostaining associated with GALT); 2 - frequent association (50-89%); and, 3 - exclusive association (immunostaining was only observed in epithelial cells overlying GALT).

Additionally, IHC was performed to characterize select T cell subsets (CD4, CD8, and Foxp3) in the cecum and colon of B6 mice (n=6 infected and control) euthanized 14 and 63 DPC. After deparaffinization and heat-induced epitope retrieval as described above, large intestinal sections were incubated with either anti-CD4 (dilution 1:250; 14-9766-82, clone 4SM95, Thermo Fisher Scientific,), anti-CD8a (dilution 1:1000; 14-0808-82, clone 4SM15, Thermo Fisher Scientific), or anti-Foxp3 (dilution 1:600; 14-5773, clone FJK-16s, Thermo Fisher Scientific) antibodies. A rabbit anti-rat secondary antibody (BA-4001, Vector Laboratories) and a polymer detection system (PK6100, Vector Laboratories) were then applied to the tissues. The 3,3’-diaminobenzidine tetrachloride (DAB) was used as the chromogen, and the sections were counterstained with hematoxylin and examined by light microscopy. Lymphoid tissues (thymus, spleen, and lymph nodes) from naïve, Cm-free, inbred mice were used as a positive control. Positive CD4, CD8, and Foxp3 immunolabeling were identified as discrete, punctate chromogenic brown dots under bright field microscopy.

In situ hybridization (ISH) - Select tissues from Cm qPCR-positive mice (n=2 B6 GI tract; n=3 S GI tract, n=3 S lungs, and n=1 S reproductive tract) were also evaluated using ISH when MOMP IHC signal was absent or inconclusive. Briefly, the target probe was designed to detect region 581-617 of *Chlamydia muridarum* str. Nigg complete sequence, NCBI Reference Sequence NC_002620.2 (1039538- C1; Advanced Cell Diagnostics, Newark, CA). The target probe was validated on reproductive tracts from mice experimentally inoculated with *Chlamydia muridarum* strain Nigg.^51^ Slides were stained on an automated stainer (Leica Bond RX, Leica Biosystems) with RNAscope 2.5 LS Assay Reagent Kit-Red (322150, Advanced Cell Diagnostics) and Bond Polymer Refine Red Detection (DS9390, Leica Biosystems). Control probes detecting a validated positive housekeeping gene (mouse peptidylprolyl isomerase B, *Ppib* to confirm adequate RNA preservation and detection; 313918, Advanced Cell Diagnostics) and a negative control, *Bacillus subtilis* dihydrodipicolinate reductase gene (dapB to confirm absence of nonspecific staining; 312038, Advanced Cell Diagnostics) were used. Positive RNA hybridization was identified as discrete, punctate chromogenic red dots under bright field microscopy.

### Flow cytometry

Following euthanasia and blood collection, the spleen was removed for immunophenotyping via flow cytometry (n=6 each experimental and control mouse/strain at 14 and 63 DPC). All remaining tissues from these mice were processed as described above. The spleen was dissociated into a single-cell suspension and RBCs lysed with ACK RBC cell lysis buffer (catalog no. A1049201, Gibco, San Diego, CA). The suspension was washed twice with PBS and blocked with Fc Receptor Block (TruStain FcX, catalog no. 101320, Biolegend, San Diego, CA) prior to staining. Cells were then stained with the following anti-mouse antibodies: APC/Fire 810 CD3 (17A2, catalog no. 100267, Biolegend), CD11b BV570 (M1/70, catalog no. 101233, Biolegend), Alexa Fluor 700 Ly-6C (HK1.4, catalog no. 128024, Biolegend), PE/Cyanine7 CD127 IL-7Rα (A7R34, catalog no. 135014, Biolegend), PerCP Ly-6G (1A8, catalog no. 127653, Biolegend), Brilliant Violet 421 CD25 (PC61, catalog no. 102034, Biolegend), Brilliant Violet 650 CD45R/B220 (RA3-6B2, catalog no. 103241, Biolegend), BUV661 NK-1.1 (PK136, catalog no. 741477, BD Biosciences, Ashland, OR), BUV496 CD4 (GK1.5, catalog no. 612952, BD Biosciences), BUV805 CD8a (5H10-1, catalog no. 752640, BD Biosciences), BUV737 F4/80 (T45-2342, catalog no. 749283, BD Biosciences), BUV395 CD45 (30-F11, catalog no. 564279, BD Biosciences), FITC CD69 (H1.2F3, catalog no. 35-0691-U100, Tonbo, Freemont, CA, PE CD44 (IM7, catalog no. 50-0441-U100, Tonbo), APC CD62L/L-selectin (MEL-14, catalog no. 20-0621-U100, Tonbo), and Live/Dead Blue (catalog no. L34961, Thermo Fisher Scientific) for discrimination of live and dead cells. Brilliant Stain Buffer (catalog no. 566349, BD Biosciences) was added to all staining cocktails. Samples were analyzed using an Aurora Spectral Analyzer (Cytek Biosciences, Fremont, CA) equipped with 5 lasers (355 nm, 405 nm, 488 nm, 561 nm, and 640 nm) with full spectral detection. Data were analyzed with FlowJo software (version 10.10, BD Life Sciences – Informatics, Ashland, OR). Population percentages were calculated for B cells, macrophages, monocytes (with separate populations of Ly6C high and low), neutrophils, NK cells, T cells, CD4+, and CD8+ cells as a subset of the total immune cells (CD45+). Additionally, the following sub-group population percentages were calculated: CD69+/CD4+, CD4+/T cells, central memory cell/CD4+, effector cells/CD4+, effector memory cells/CD4+, naïve cells/CD4+, T regulatory cells/CD4+, CD8+/T cells, CD69+/CD8+, central memory/CD8+, effector/CD8+, effector memory/CD8+ and naïve/CD8+. The gating strategy to detect myeloid and lymphoid cell populations is shown in Figure 7A.

Additionally, following euthanasia and blood collection, the gastrointestinal tract (from duodenum to colon; B6 mice only) was removed for immunophenotyping via flow cytometry (n=6 each experimental and control at 14 and 63 DPC). The duodenum, jejunum, and ileum were separated from the cecum and colon, with the former pooled as one sample for small intestine and the latter for large intestine. The small and large intestines were each dissociated into single-cell suspensions, and RBCs were lysed with ACK RBC cell lysis buffer (catalog no. A1049201, Gibco). Half of each sample was stimulated with phorbol 12-myristate 13-acetate (PMA, 50 ng/mL), ionomycin (750 ng/mL) + brefeldin A (BFA, 10 ug/mL; all obtained from Sigma-Aldrich, St Louis, MO) for 3.5hrs for cytokine induction prior to staining; the other half was kept as a single cell suspension for use in cell population analysis. The suspension was washed twice with PBS and blocked with Mouse BD Fc Block (2.4G2, catalog no. 553141, BD Biosciences) prior to staining. Cells were then stained with the following anti-mouse antibodies: Brilliant Violet 785 CD45 (30-F11, catalog no. 103149, Biolegend), PerCP/Cyanine 5.5 CD5 (53-7.3, catalog no. 100624, Biolegend), Alexa Fluor 488 TCR γ/δ (GL3, catalog no. 118128, Biolegend), APC/Fire 750 CD11b (M1/70, catalog no. 101262, Biolegend), APC/Fire 770 CD11c (N418, catalog no. 117352, Biolegend), APC/Fire 750 CD45R/B220 (RA3-6B2, catalog no. 103260, Biolegend), APC/Cyanine7 CD19 (6D5, catalog no. 115530, Biolegend), Alexa Fluor 700 CD90.2/Thy 1.2 (30-H12, catalog no. 105320, Biolegend), Brilliant Violet 711 CD127/IL-7Rα (A7R34, catalog no. 135035, Biolegend), Brilliant Violet 605 CD196/CCR6 (29-2L17, catalog no. 129819, Biolegend), CD335/NKp46 (29A1.4, catalog no. 137604, Biolegend), PE/Cyanine7 Ly-6G (1A8, catalog no. 127618, Biolegend), PE/Cyanine7 IFNγ (XMG1.2, catalog no. 505826, Biolegend), CD3e (145-2C11, catalog no. 45-0031-82, Thermo Fisher Scientific), NK1.1 (PK136, catalog no. 45-5941-82, Thermo Fisher Scientific), retinoid-related orphan receptor (RORγ; B2D, catalog no. 61-6981-82, Thermo Fisher Scientific), T-bet (4B10, catalog no. 50-5825-82, Thermo Fisher Scientific), FOXP3 (FJK-16s, catalog no. 14-5773-82, Thermo Fisher Scientific), TNF alpha (MP6-XT22, catalog no. 48-7321-82, Thermo Fisher Scientific), BUV395 CD4 (GK1.5, catalog no. 563790, BD Biosciences), BUV395 GATA3 (L50-823, catalog no. 565448, BD Biosciences), BUV395 IL-17A (TC11-18H10, catalog no. 565246, BD Biosciences), and Ghost Dye Violet 510 (catalog no. 13-0870-T500, Tonbo) for discrimination of live and dead cells. Samples were analyzed with a BD Foressa II flow cytometer (BD Biosciences, San Jose, CA) equipped with 5 lasers (355 nm, 405 nm, 488 nm, 561 nm and 640 nm) and the software above. Population percentages were calculated for cells (and subpopulations) including: CD4^+^ T effector cells (Foxp3^-^ and RORγ^+^ or T-bet^+^), CD4^+^ T regulatory cells (Foxp3^+^ and GATA3^+^ or RORγ^+^), innate lymphoid cells (T-bet^+^ for ILC1, GATA3^+^ for ILC2, and RORγ^+^ for ILC3), and ILC3 subsets (T-bet^+^ or CCR6^+^ among RORγ^+^ ILC3s). Additionally, intracellular cytokine expression was assessed in CD4^+^ T cells (IFNγ, IL-17A, IL-22, TNF) and ILC3s (GMCSF, IL-17A, IL-22, TNF) populations for both small and large intestines after cell fixation and permeabilization. The gating strategy to detect innate lymphoid cell (ILC), myeloid, and lymphoid cell populations is shown in Figure 9A.

### Cytokine secretion assays

Select cytokines, chemokines, or growth factors associated with inflammatory responses were measured in sera from B6 mice (n=6 experimental; n=2 control at 14 and 63 DPC). Briefly, blood from Cm-infected or Cm-free control mice were collected at necropsy and centrifuged at high speed to separate serum. Sera (80 µl) from each mouse diluted 1:1 with sterile PBS was used to measure granulocyte-macrophage colony stimulating factor (GM-CSF), IFNγ, IL-1β, IL-2, IL-4, IL-6, IL-10, IL-12p70, monocyte chemoattractant protein 1 (MCP-1), and tumor necrosis factor alpha (TNFα) using Luminex XMAP technology (Mouse Focused 10-Plex Discovery Assay; Eve Technologies, Calgary, CN). Assay sensitivities ranged from 0.4 – 10.9 pg/mL for the analytes.

### Statistical analysis

Due to the lack of data normality, Mann-Whitney U (Wilcoxon Rank Sum) tests were used to determined statistical differences in hemogram, leukogram, and serum biochemistry analyte concentrations between infected and control mice for each strain/stock. Mann Whitney U tests were also used to compare mean GALT hyperplasia scores between infected and control mice for each strain/stock at 14 and 63 DPC. Kruskal-Wallis tests (B6 vs. C vs. S) or Mann-Whitney U tests (B6 vs. C; B6 vs. S; C vs. S) were performed to assess differences in the localization of Cm MOMP antigen in epithelial cells overlying GALT at each timepoint. Flow cytometry data was analyzed as the proportion of each specific immune cell type to its parent population (e.g., either total immune cells, CD4+ T cells, ILCs, etc.); Mann-Whitney U tests were used to determine statistical differences between the experimental and control groups of the 3 mouse strains/stocks at each timepoint. Mann-Whitney U tests were also utilized to find significant systemic differences in serum cytokine values between infected and control B6 mice at 14 and 63 DPC. For all tests, statistical significance was set as a *P* value of equal to or less than 0.05. Analyses were performed in JMP Pro (version 17, SAS Institute, Cary, NC). Graphical representations were created using Prism 9 for Windows (version 9.3.0, GraphPad Software, Boston, MA).

## Results

### Clinical impact of Cm infection

No experimentally inoculated or cohoused Cm-infected mice developed clinical signs of illness throughout duration of the experiment.

### Fecal shedding of Cm

All fecal samples from experimentally inoculated mice were Cm positive by qPCR (Table 1). These mice were positive at the first collection point 7 DPI (82,202 +/- 16,350, mean copy number +/- standard error of the mean [SEM]) and continued shedding through 30 (90,774 +/- 33,560) and 75 (11,165 +/- 2,766) DPI. Just prior to cohousing (at 95 DPI), inoculated C mice remained positive with reduced Cm fecal shedding (3,192 +/- 649). The highest individual Cm copy number was observed at 30 DPC (214,453). All control mice remained Cm qPCR negative.

**Table 1.** Cm 23S rRNA copy number in pooled fecal samples from C mice infected via orogastric gavage.

| Day | Copy #<sup>a</sup> |  |  |  |
| --- | --- | --- | --- | --- |
|  | Min | Max | Mean | SEM |
| 7 | 13,159 | 106,734 | 82,203 | 16,350 |
| 30 | 26,439 | 214,453 | 90,774 | 33,560 |
| 75 | 1,622 | 53,122 | 11,165 | 2,766 |
| 95 | 402 | 13,159 | 3,192 | 649 |

<sup>a</sup> Copy number calculated from Ct values of qPCR assays

Results for all cohoused mice are displayed graphically in Figure 2A. All B6 mice were Cm positive by fecal qPCR as early as 3 DPC (100 +/- 50) and remained colonized through the end of the study at 180 DPC (75,542 +/- 31,191). The highest copy numbers were seen between 7 (601,926 +/- 568,902) and 49 DPC (567,281 +/- 184,588). One of 3 C mice was Cm positive at 3 DPC (402); by 7 DPC all C mice were shedding Cm (152,576 +/- 139,358). The highest average copy numbers were seen between 30 (350,348 +/- 173,667) and 49 DPC (228,228 +/- 77,520) with a single C mouse shedding a large amount of at 7 DPC (430,887). C mice shed Cm 180 DPC (35,333 +/- 8,894). No S mice were shedding at either 3 or 7 DPC, and only 1of 6 S mice was shedding at 14 DPC (200). Despite this delay in shedding, all S mice were shedding at 30 DPC with the highest copy numbers measured between 30 (349,848 +/- 81,038) and 49 DPC (975,472 +/- 274,527). S mice shed through 180 DPC (15,382 +/-5,848). Despite temporal differences in time to initial shedding between strains and stocks, peak copy numbers were similar. All control mice remained Cm qPCR negative. An error in sample collection resulted in feces collected from all mice at 21 DPC to be formalin fixed invalidating qPCR results for the timepoint.

**Figure 2.**
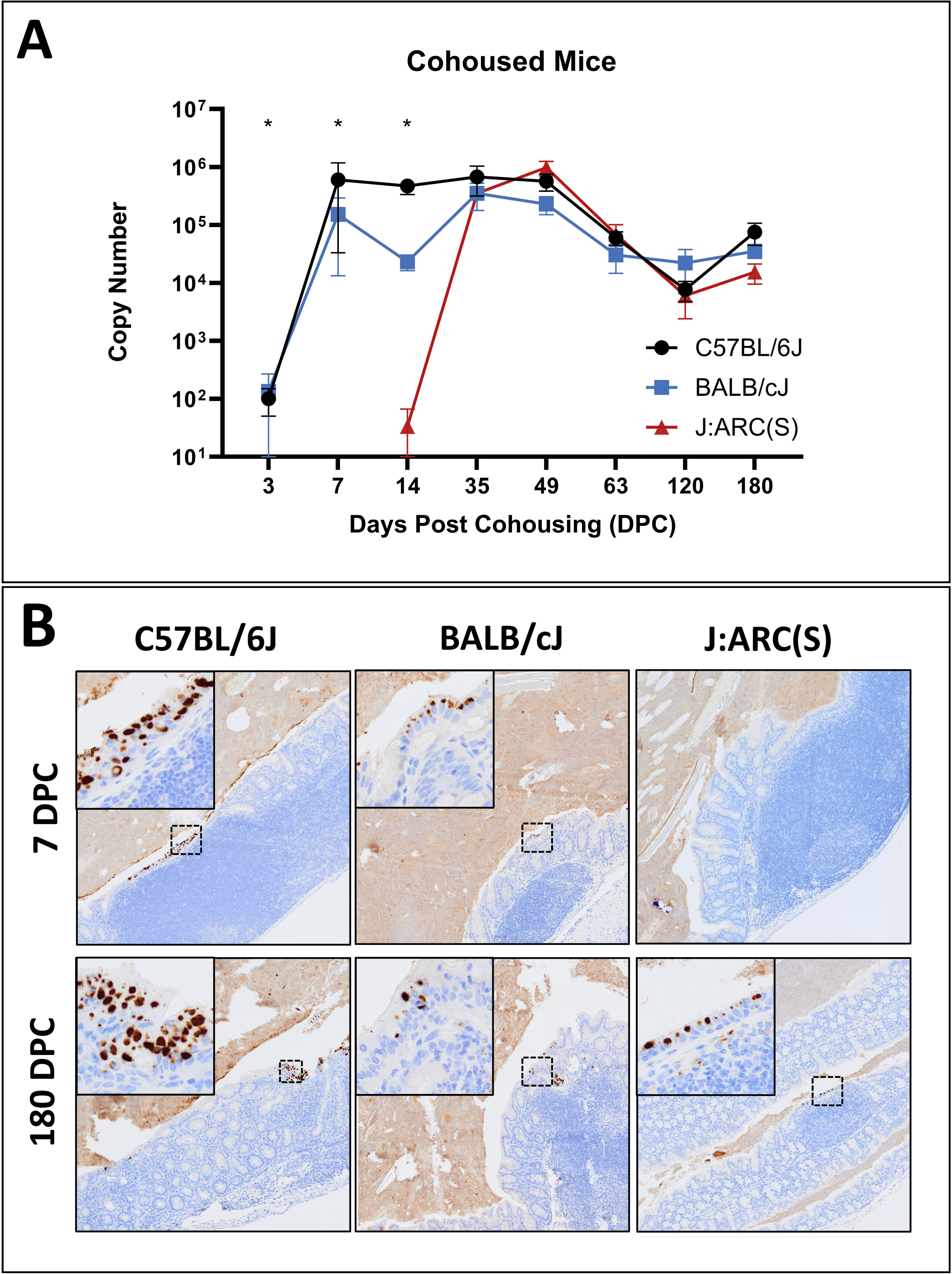
Fecal shedding and intestinal colonization with Cm.

(A) Fecal qPCR data (mean Cm copy number +/- SEM) by timepoint and strain/stock (n=3-6 mice/timepoint).

*= significant difference between C57BL/6J (B6) and BALB/cJ (C), and the J:ARC(S) [S] strain/stock (Kruskal-Wallis test, p≤0.05).

(B) Immunohistochemical staining for Cm MOMP. Cm antigen immunolabeling in surface epithelial cells throughout the colon (intracellular brown staining) of B6, C and S mice at 7 and 180 DPC. Cm inclusions are noted in the cytoplasm of enterocytes (insets). Immunolabeling is detected in B6 and C mice at 7 DPC and all strains/stocks at 180 DPC. Staining of luminal contents reflects non-specific staining of extracellular debris Immunohistochemistry for MOMP antigen, 2x (insets 10x).

### Clinical and anatomic pathology

There were no clinically or statistically significant changes in the hemograms, leukograms, or serum biochemistry analyte concentrations between Cm-infected and control mice of any strain/stock, nor were there consistent or replicable patterns of temporal change (data not shown).

Gross lesions were not appreciated at necropsy in Cm-inoculated, cohoused, or control mice. Microscopic lesions in both Cm-inoculated and Cm-infected cohoused mice were limited to the respiratory (cohoused only) and the GI tracts (both). There were no significant histologic lesions associated with Cm in the lungs or reproductive tract in any mouse evaluated at any timepoint throughout the study. One infected B6 mouse at 7 DPC had minimal neutrophilic inflammation associated with Cm IBs in the nasopharynx. The lungs of Cm-infected and control mice at various time points post-cohousing had mild, focal to multifocal, peribronchiolar and perivascular lymphocytic and histiocytic infiltrates consistent with spontaneous background lesions.^19,50^ Additional incidental pulmonary findings included acidophilic macrophagic pneumonia (1/6 infected B6 at 14 DPC; 1/1 control B6 and 1/3 infected C at 120 DPC) and occasional mild, follicular and germinal center hyperplasia of the tracheobronchial lymph nodes (2/6 infected C at 63 DPC; 1/3 infected and 1/1 control B6, and 1/3 infected and 1/1 control S mice at 180 DPC). There were no IBs detected in the trachea or lungs in any mouse at any timepoint.

Histological analysis of the gastrointestinal tract showed that Cm IBs were occasionally detected in the cytoplasm of epithelial cells throughout the intestines in Cm qPCR+ mice of all strains/stocks (both inoculated and cohoused). IBs were most frequently observed in the cecum and colon, however they were occasionally observed in the small intestine. Additionally, minimal-to-mild enteritis, typhlitis, and/or colitis were occasionally observed. The majority of enteritides/colitides were observed in Cm-infected mice (15/18 mice with GI tract inflammation), however uninfected control mice (3/18) were also affected. These inflammatory changes were most commonly observed in B6 (10/18) and S (7/18) mice. Typically, the lamina propria was multifocally infiltrated by low numbers of mixed lymphocytes, plasma cells, and/or histiocytes which occasionally infiltrated the submucosa (Figure 3). Cm IBs were found with and without associated inflammation (Figure 3). Mild crypt cell hyperplasia (6/18) and crypt abscessation (1/18) were also infrequently observed.

**Figure 3.**
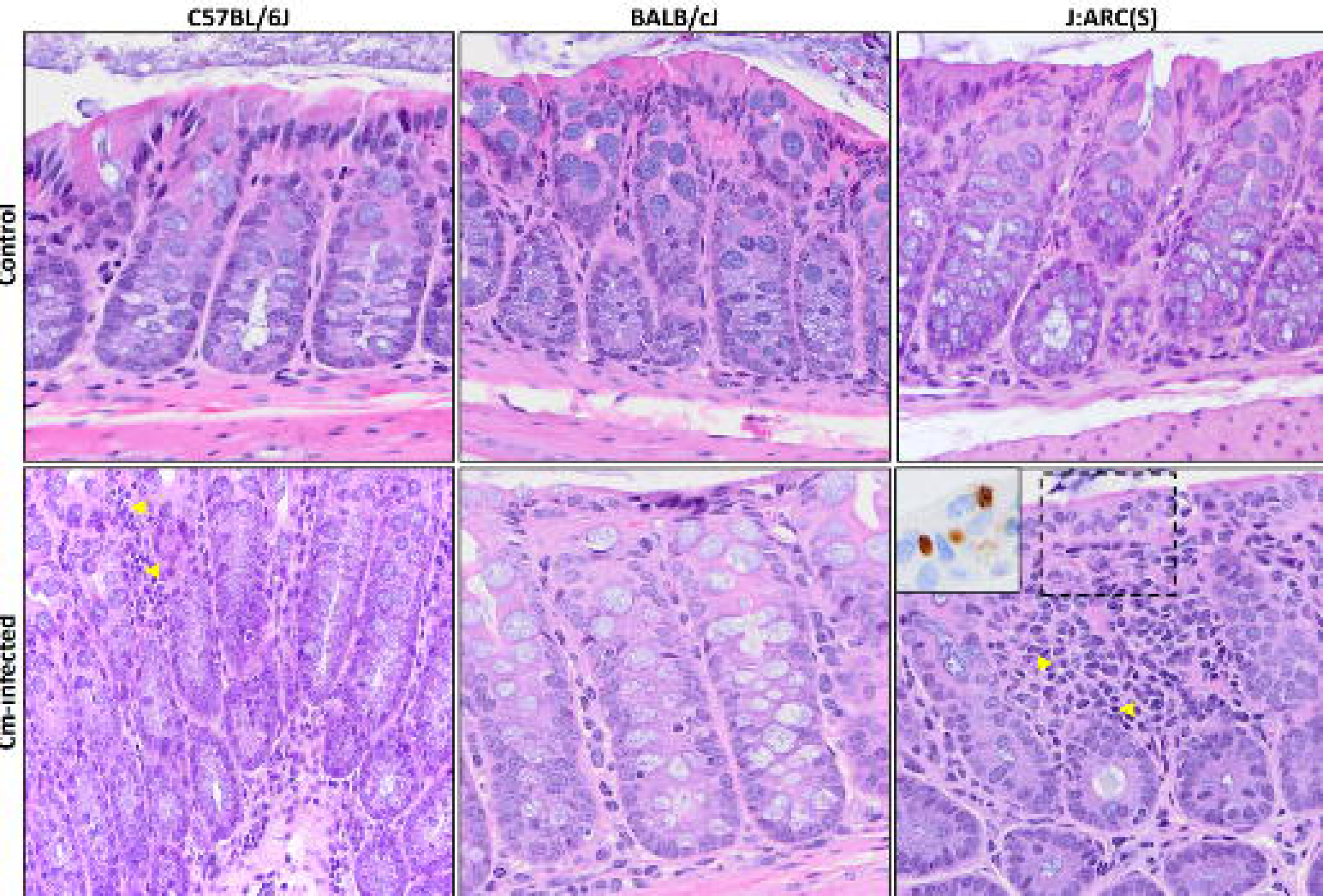
A subset of Cm-infected mice exhibited mild colitis at 180 DPC. (top row) Representative histology of the colon from uninfected (control) C57BL/6J (B6), BALB/cJ (C), and J:ARC(S) [S] mice demonstrating normal intestinal mucosa and submucosa. H&E, 60x. (bottom row) A subset of Cm- infected B6 and S mice developed mild multifocal colitis at 180 DPC. The lamina propria and/or submucosa was infiltrated by lymphocytes, plasma cells, (yellow arrowheads), histiocytes, and less frequently neutrophils. No colitis was observed in Cm-infected C mice. Inset depicts Cm MOMP immunolabeling (dashed box) in the cytoplasm of colonic enterocytes from infected C mice. H&E, 60x (inset: immunohistochemistry for Cm MOMP antigen).

Cm-infected mice exhibited GALT hyperplasia which was more extensive in distribution and frequency as compared to uninfected control mice of the same strain at the same timepoint (days 14 and 63 DPC depicted in Figure 4A). Given the small sample sizes (n=3-6 infected and 1-6 control/timepoint), differences were not statistically significant; however, several notable trends were observed. Hyperplasia was not detected in any mice at 3 or 7 DPC. With a single exception (below), GALT hyperplasia was detected in the majority of Cm-infected cohoused mice in all 3 strains/stock thereafter (14 through 180 DPC). At each of these timepoints, GALT hyperplasia was either absent, detected in a minority, and/or detected with lower scores in uninfected counterparts. At 21 DPC, GALT hyperplasia was not observed in any of the Cm-infected S mice assessed.

**Figure 4.**
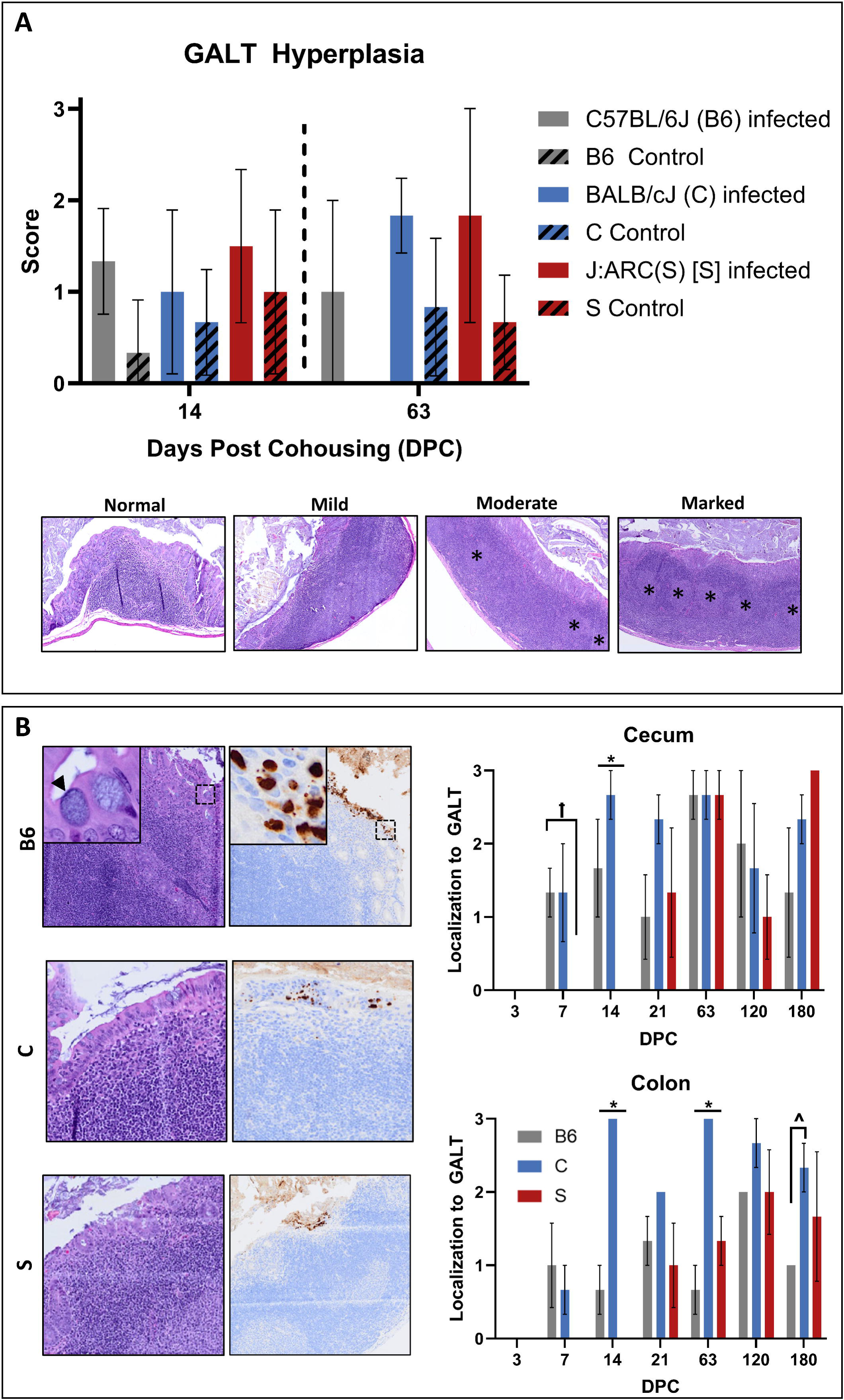
Cm large intestinal colonization is associated with GALT. (A) GALT hyperplasia scores (mean +/- SEM) at 14 and 63 DPC in C57BL/6J (B6, n=3/timepoint), BALB/cJ (C, n=6/timepoint), and J:ARC(S) [S, n=6/timepoint] mice. No significant differences in GALT hyperplasia were appreciated due to a small sample size, however infected mice had higher scores at both 14 and 63 DPC. Representative examples of normal (0), mild (1), moderate (2), and marked (3) GALT hyperplasia in the large intestine from S mice at 63 DPC are presented. * = prominent germinal centers. H & E, 5x. (B) Association between Cm-infected epithelial cells and GALT in B6, C, and S mice (n=3-6 mice/timepoint). Cm infection of epithelial cells overlying GALT provided in images on left. Arrowhead=IB; Cm MOMP antigen (brown staining) present in epithelium overlying GALT in photomicrographs. IHC for MOMP. 2x (insets 20x). GALT association scores (mean +/- SEM) in the cecum and colon of B6, C, and S mice at various times post-cohousing in the histogram on the right. * = significant differences (p=<0.05) between each strain/stock; ^= significant differences (p=<0.05) between B6 and C mice; [ = significant differences (p=<0.05) between B6 and S mice.

Strong, specific Cm MOMP antigen immunolabeling was observed almost exclusively in the cytoplasm of surface epithelial cells throughout the small and large intestines in both Cm-inoculated and Cm-infected cohoused mice (Figure 2B). Immunolabeling was present at all timepoints where mice were shown to be shedding Cm in their feces, e.g., B6 and C mice at both 7 and 180 DPC; S mice at 180 DPC, and absent when fecal qPCR was negative, e.g., S mice at 7 DPC (Table 2). This immunolabeling was detected primarily throughout the cecum and colon, however occasional signal was detected in the small intestine, and a single C mouse demonstrated immunostaining in the surface epithelial cells of the gastric limiting ridge at 14 DPC. The B6 mouse with the observed neutrophilic inflammation in the nasopharynx had a single Cm MOMP antigen staining nasal epithelial cell. There was no signal detected in the lungs nor the reproductive tissues of any infected cohoused mouse nor was there immunostaining in any tissue at any timepoint in the uninfected control mice.

**Table 2.** Cm fecal qPCR and MOMP antigen IHC results.

| Day | C57BL/6J<sup>a</sup> |  |  | BALB/cJ<sup>a</sup> |  |  | J:ARC(S)<sup>a</sup> |  |  |
| --- | --- | --- | --- | --- | --- | --- | --- | --- | --- |
|  | PCR | SI IHC | LI IHC | PCR | SI IHC | LI IHC | PCR | SI IHC | LI IHC |
| 3 | 3/3 | 0/3 | 0/3 | 1/3 | 1/3 | 1/3 | 0/3 | 0/3 | 0/3 |
| 7 | 3/3 | 2/3 | 3/3 | 3/3 | 1/3 | 3/3 | 0/3 | 0/3 | 0/3 |
| **14** | 6/6 | 3/3 | 3/3 | 6/6 | 4/6 | 6/6 | 1/6 | 0/6 | 1/6 |
| **21** | n/a | 2/3 | 3/3 | n/a | 1/3 | 3/3 | n/a | 1/3 | 3/3 |
| **63** | 6/6 | 2/3 | 3/3 | 6/6 | 2/6 | 6/6 | 6/6 | 1/6 | 6/6 |
| **120** | 3/3 | 0/3 | 3/3 | 3/3 | 0/3 | 3/3 | 3/3 | 0/3 | 3/3 |
| **180** | 3/3 | 0/3 | 3/3 | 3/3 | 1/3 | 3/3 | 3/3 | 0/3 | 3/3 |

<sup>a</sup> Data presented as number positive in the total number sampled at that timepoint. n/a= not available

Cm MOMP antigen immunolabeling was frequently observed in cecal and colonic epithelia overlying GALT in Cm-infected cohoused mice. This association was not seen in the single C mouse with Cm MOMP immunostaining at 3 DPC; no other C and all B6 and S mice had immunostaining at this timepoint. At 7 DPC, B6 presented with higher GALT association scores in the cecum (1.33 +/- 0.33; mean +/- SEM) than S mice (0; no immunostaining observed). Additionally, B6 and C mice had higher cecal (B6 1.67 +/- 0.67; C 2.67 +/- 0.21) and colonic (B6 0.67 +/- 0.33; C 2.83 +/- 0.17) scores as compared to S mice (0; no immunostaining observed) at 14 DPC. At 63 DPC, C mice presented with significantly higher colonic scores (2.83 +/- 0.17) than both B6 (0.67 +/- 0.33) and S mice (1.3 +/- 0.33). Again at 180 DPC, C mice presented with significantly higher association scores (2.33 +/- 0.33) in the colon than B6 (entire cohort 1.0) and S mice (1.67 +/- 0.88). All these differences were statistically significant (p≤0.05). Cm MOMP immunolabeling was occasionally noted in large intestinal surface epithelial cells which was not GALT associated. Cm MOMP staining in the SI was observed in markedly fewer cells, was of lesser or equal intensity, and was only rarely GALT associated. There was no Cm MOMP antigen detected in any of the control mice. The presence of IBs in the cytoplasm of surface epithelial cells throughout the gastrointestinal tract was correlated with Cm MOMP antigen immunolabeling (Figure 4B).

The T cell composition of the GALT evaluated in B6 mice at 14 and 63 DPC consisted of both CD4^+^ and CD8^+^ T cells. Both cell types were present throughout the small and large intestine. In the GALT from infected mice, CD4^+^ and CD8^+^ T cells frequently clustered in interfollicular regions between germinal centers and were also scattered in the perifollicular regions and mantle zones. CD4^+^ T cells were frequently detected in the GALT from both anatomic sites, with appreciably greater CD4 T cell immunolabeling in the large intestine. Conversely, control mice demonstrated a more balanced T cell composition, with roughly equal staining in area and intensity of both CD4^+^ and CD8^+^ T cell populations. There was minimal Foxp3 signal throughout the lamina propria and GALT in the small and large intestines of both infected and control mice. All cell types assessed were located in the inter- and perifollicular GALT regions, consistent with anatomic location of T cells in murine GALT. Examples are shown in Figure 5.

**Figure 5:**
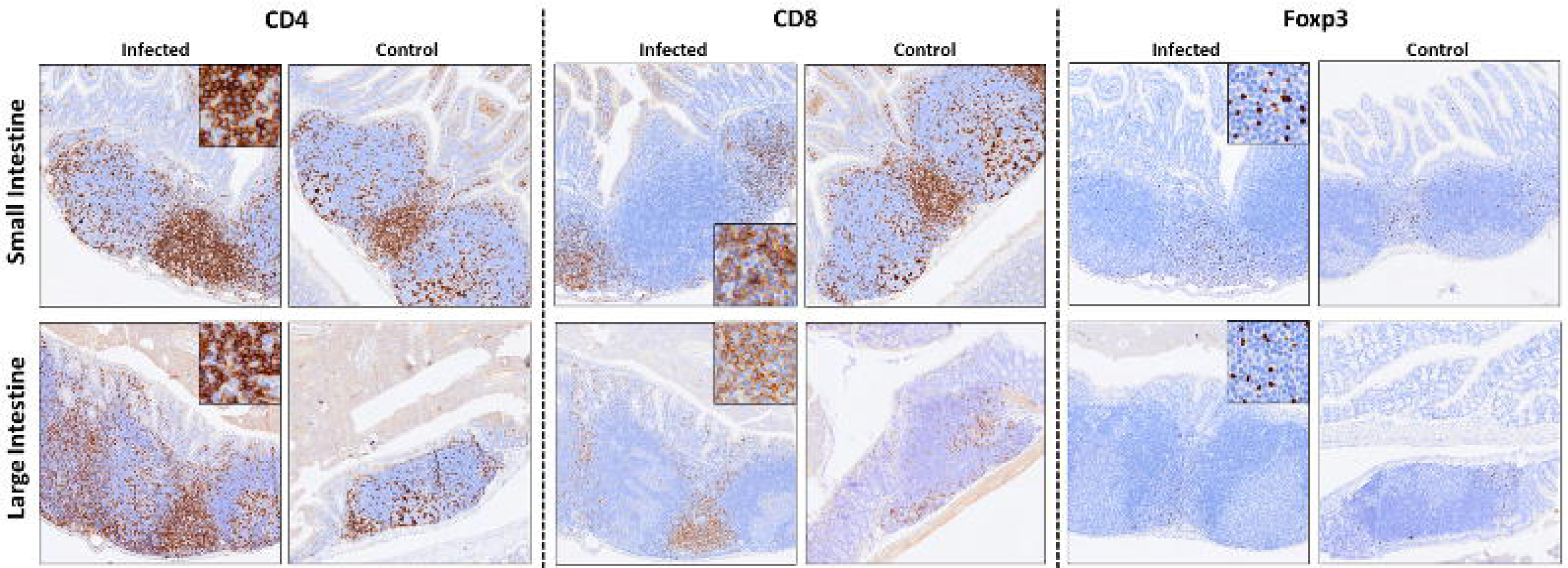
T cell subsets in small and large intestinal GALT. Representative immunohistochemistry for CD4, CD8 and Foxp3 cells in the GALT of C57BL/6J mice at 14 DPC. Infected mice had T cell populations comprised of primarily CD4 positive cells (left panels, brown cytoplasmic/membranous immunolabeling of round cells) with less frequently observed CD8 positive cells (middle panels, brown cytoplasmic/membranous immunolabeling of round cells). Both CD4 and CD8 positive cells were often found clustering in interfollicular regions between germinal centers and scattered in the mantle zones, perifollicular regions, and associated lamina propria. The small intestine from the uninfected control mice had similar immunolabeling of CD4 and CD8 positive lymphocytes as found in infected mice, but the CD4 positive cells were less obvious in controls. The lymphocyte populations of CD4 and CD8 cells in the large intestine were increased and more homogenous in uninfected controls. Both infected and uninfected mice presented with infrequent T regulatory cells (assessed via Foxp3; right panels, brown nuclear immunolabeling). Immunohistochemistry for CD4, CD8, and Foxp3, 5x (insets 20x).

Cm nucleic acid was detected via ISH in all intestinal tissues where IHC signal was either absent or inconclusive in Cm fecal qPCR+ mice (n=2 B6 and 3 S mice). No nucleic acid was detected with ISH in the pulmonary (n=3 S mice) or reproductive (n=1 S mouse) tissues evaluated. Examples are provided in Figure 6.

**Figure 6:**
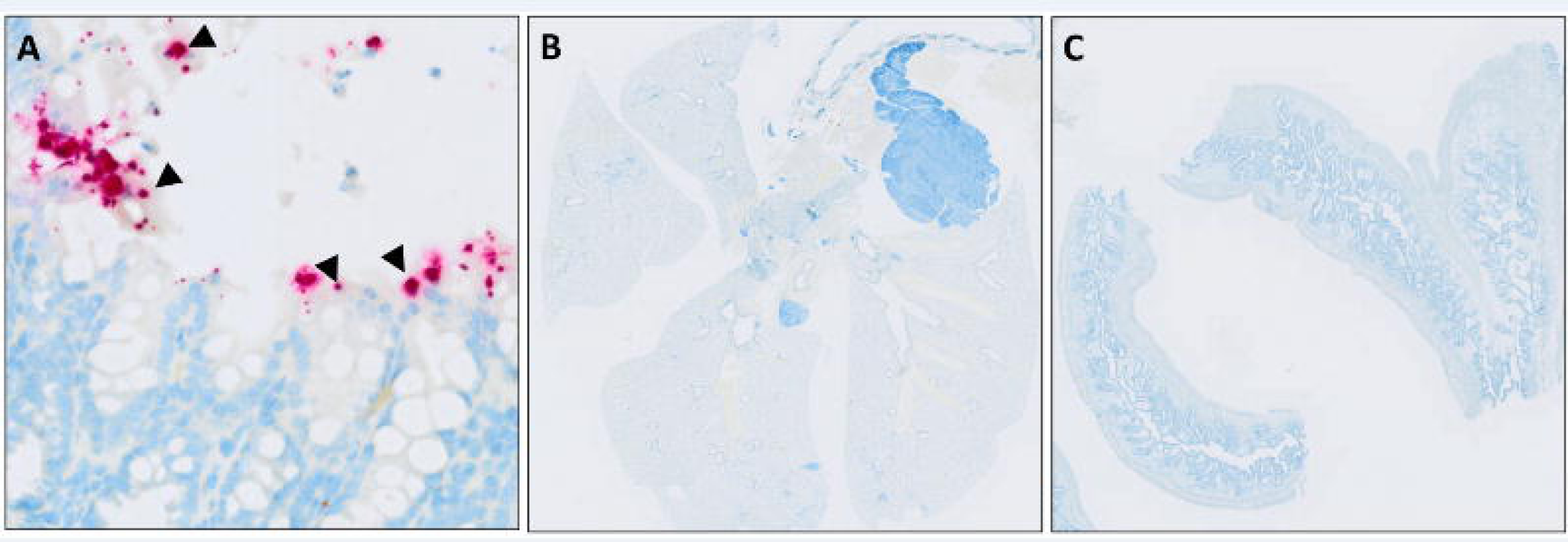
ISH staining for Cm in the gastrointestinal tract, lungs and uterus. (A) Cm nucleic acid (punctate dot red staining, arrowheads) is detected in apical epithelial cells of the colon in a Cm-infected C57BL/6J mouse, 63 DPC. ISH for Cm, 10x. (B) Lungs from an Cm-infected J:ARC(S) [S] mouse, 63 DPC demonstrating no Cm nucleic acid hybridization. ISH for Cm, subgross. (C) Cm nucleic acid is not detected in the uterus from a Cm-infected S mouse at 14 DPC. ISH for Cm, subgross.

### Cm in male mice

All Cm-inoculated male C, B6, and S mice were confirmed to be shedding and remained asymptomatic throughout the experiment. IBs were visualized in surface epithelial cells in association with strong, specific Cm MOMP immunolabelling throughout the cecum and colon (data not shown); no other histologic lesions nor Cm MOMP immunolabeling were appreciated in any other tissues including the reproductive tract. Accordingly, there were no clinical, histologic, or immunohistochemical differences appreciated between Cm-infected male and female mice.

### Immunophenotyping

Limited statistically significant differences were detected in splenic cell populations between infected and uninfected mice of each strain/stock. All results described were significant at p≤0.05 unless otherwise noted. There were few consistent changes detected among strains/stocks other than a monocytosis at 63 DPC. While there were shifts in T-cell population subtypes, the changes differed by strain/stock. Results for B6 are provided in Figure 7B. Results for the C and S mice are provided in Figure 8.

**Figure 7.**
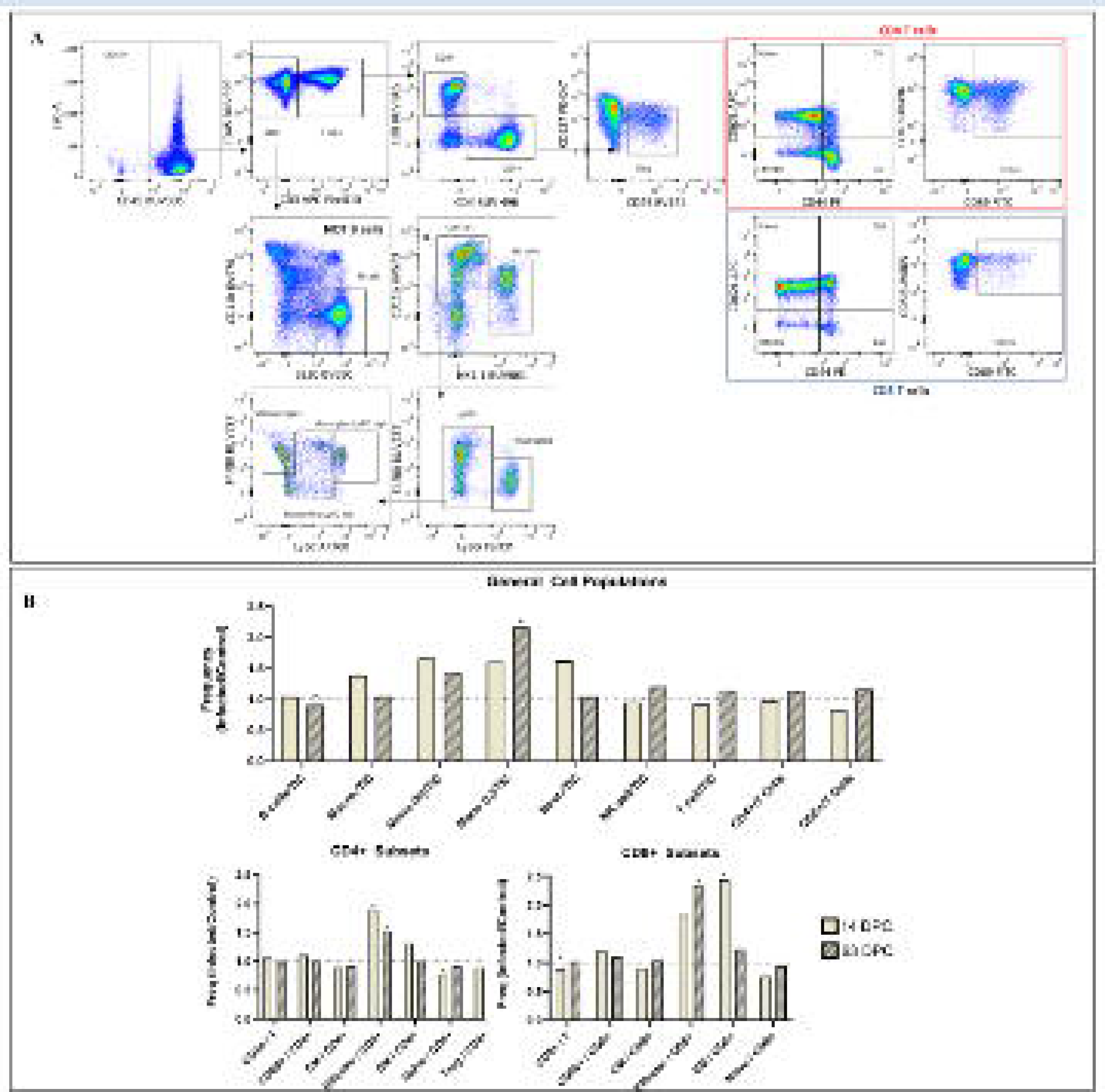
Immunophenotyping of splenic immune cells during early and late stages of Cm infection in C57BL/6J (B6) mice. (A) Gating strategy defining the immune subpopulations of splenic cells after excluding debris, doublets, and dead cells. Splenic immune cells were first identified as CD45^+^ cells. B cells were then quantified as CD3^-^/B220^+^. T cells were identified as CD3^+^. CD8 and CD4 T cells were identified by expression of their respective marker, while the subpopulation of T_reg_ was identified as CD4^+^/CD25^+^/CD127^-^. CD4^+^ and CD8^+^ T cell activation was identified as CD69^+^. Neutrophils and monocytes were quantified as CD11b+, and further distinguished as Ly6G+ and Ly6G–Ly6C+, respectively. Data displayed are from B6 mice at 14 DPC. (B) Infected B6 mice demonstrated increased frequencies of myeloid and lymphoid cell population subtypes when compared to uninfected controls, including monocytes (63 DPC), effector CD4+ T cells (14 and 63 DPC), effector CD8+ T cells (63 DPC) and effector memory CD8+ T cells (14 DPC). Other population increases did not reach statistical significance. Additionally, infected B6 mice had fewer B cells (63 DPC), naïve CD4+ cells (14 DPC), and overall CD8+ T cells (14 FPC). Data represent the frequencies of specific cell populations as compared with the parent population and are displayed as the proportions of average values in infected as compared to control mice (n=6 mice/group). Dotted line (1.0) indicates no difference in frequency of detection between control and infected mice. Values above this line represent an increased frequency of detection in infected mice and values below this line represent a decreased frequency of detection in infected mice. macro = macrophage; mono = monocyte; neut = neutrophil; EM = effector memory cell; TIC = total immune cells. *=p≤0.05.

**Figure 8.**
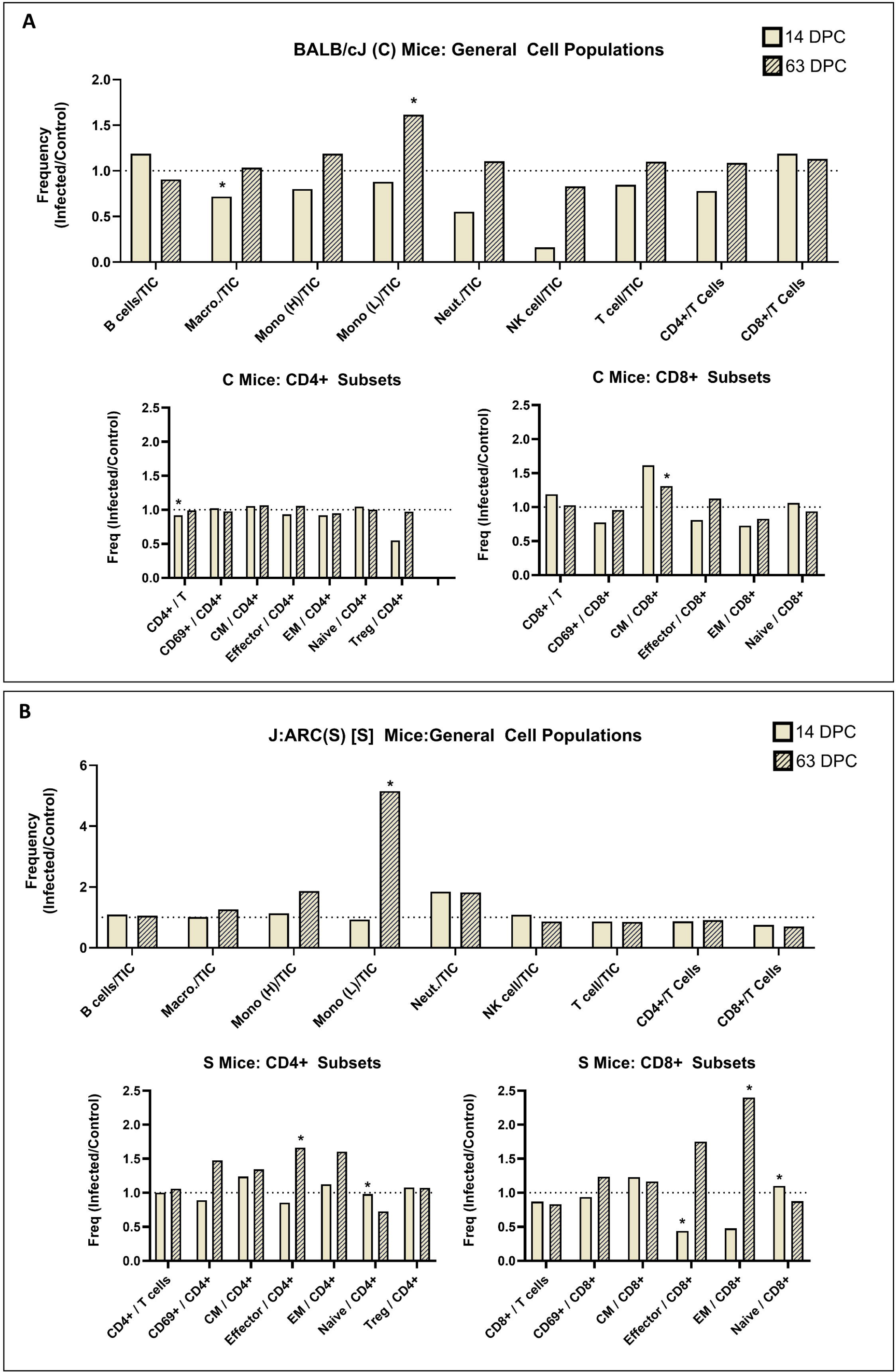
Immunophenotyping of splenic immune cells during early and late stages of Cm infection in BALB/cJ (C) and J:ARC(S) [S] mice. Data (A and B) represent the frequencies of specific cell populations as compared with the parent population and are displayed as proportions of average values of infected as compared to control mice (n=6 mice/group). Dotted line (1.0) indicates no difference in frequency of detection between control and infected mice. Values above this line represent an increased frequency of detection in infected mice and values below this line represent a decreased frequency of detection in infected mice. (A) Infected C mice demonstrated increased frequencies of myeloid and lymphoid cell population subtypes when compared to uninfected controls, including monocytes (63 DPC) and central memory CD8+ T cells (63 DPC). Additionally, Cm-infected mice demonstrated fewer macrophages and CD4+ T cells when compared to uninfected controls at 14 DPC. (B) Infected S mice demonstrated increased frequencies of myeloid and lymphoid cell population subtypes when compared to uninfected controls, including monocytes (63 DPC), effector CD4+ T cells (63 DPC), effector memory CD8+ T cells (63 DPC), and naïve CD8+ T cells (14 DPC). Additionally, infected mice demonstrated fewer naïve CD4+ T cells and effector CD8+ T cells when compared to uninfected controls at 14 DPC. macro = macrophage; mono = monocyte; neut = neutrophil; EM = effector memory cell; TIC = total immune cells. * =p≤0.05.

Infected B6 mice demonstrated a decrease in B cells (CD3^-^, B220^+^) and an increase in monocytes (CD11b^+^, Ly6C^+^) at 63 DPC. CD4^+^ T cells in infected B6 mice were more frequently effector cells (T_eff_; CD4^+^, CD62L and CD44^-^) at both 14 and 63 DPC, or effector memory cells (CD4 and CD44^+^, CD62L^-^) and less frequently naïve (CD4 and CD62L^+^, CD44^-^) T cells at 14 DPC. Differences were also noted in the CD8^+^ T cell population with Cm-infected B6 mice demonstrating fewer CD8^+^ cells, but more frequent effector memory cells (CD8 and CD44^+^, CD62L^-^) at 14 DPC. However, there were greater numbers of effector cells (CD8^+^, CD62L and CD44^-^) at 63 DPC.

Infected C mice had fewer macrophages (CD11b and F4/80^+^, Ly6C^-^) at 14 DPC while they had greater numbers of monocytes present at 63 DPC. There were limited differences in the T cell subsets in infected C mice; they had fewer CD4^+^ T cells at 14 DPC and more central memory CD8^+^ T cells (CD8, CD44, and CD62L^+^) at 63 DPC.

Infected S mice demonstrated a marked monocytosis at 63 DPC. T cell populations in infected S mice were characterized by decreased effector cells and increased naïve CD8^+^ T cells at 14 DPC, and increased central memory cells (CD4, CD44, and CD62L^+^), decreased naïve CD4^+^ T cells and increased effector memory cells at 63 DPC.

Immunophenotyping of the gastrointestinal tract of B6 mice revealed a prolonged local inflammatory response characterized by an increase in myeloid cells, and a shift toward effector and away from memory cells with cells expressing both Th1 and Th17 cell response markers. Infected mice demonstrated sustained increases in myeloid cells (Ly6G^+^). Both effector T cells and ILC3s demonstrated a sustained elevation of cytokine production. There were greater numbers of immune cells (CD45^+^) in the large intestine of infected mice as compared to uninfected controls at both 14 and 63 DPC. Of the CD4^+^ T cells present, Th1 cells (CD4 and T-box transcription factor [T-bet]^+^, Foxp3^-^) were more frequently observed. The T_eff_ cells exhibited greater T-bet and RORγt expression at 63 DPC, consistent with Th1 and Th17 responses. While there were similar trends observed at 14 DPC, the differences were not significant. There were fewer regulatory T cells (T_reg_; CD4 and Foxp3^+^) present at both 14 and 63 DPC. While there were not significant differences appreciated in T_reg_ subsets, they were more frequently both GATA3 and RORγ negative. Infected mice also demonstrated an increase in T-bet^+^ ILCs at 63 DPC (CD90, CD127, and Tbet^+^) with a similar but not a significant trend at 14 DPC. Greater T-bet^+^ and CCR6^+^ ILC3s cells (CD90, CD127, Tbet, CCR6^+^) were found in infected mice at 63 DPC. CD4^+^ cells from infected mice expressed more cytokines including IFNγ, IL-17A, and IL-22 at both 14 and 63 DPC. ILC3 cell populations expressed greater amounts of cytokines including IL-17A at 14 DPC, GMCSF at 63 DPC, and IL-22 at both 14 and 63 DPC. Small intestinal changes were limited. The number of total immune cells did not increase significantly following infection; however, the ILC3 cells of infected mice more frequently expressed CCR6 at 14 DPC and T-bet at both 14 and 63 DPC. Results for large intestine are provided in Figure 9B.

**Figure 9.**
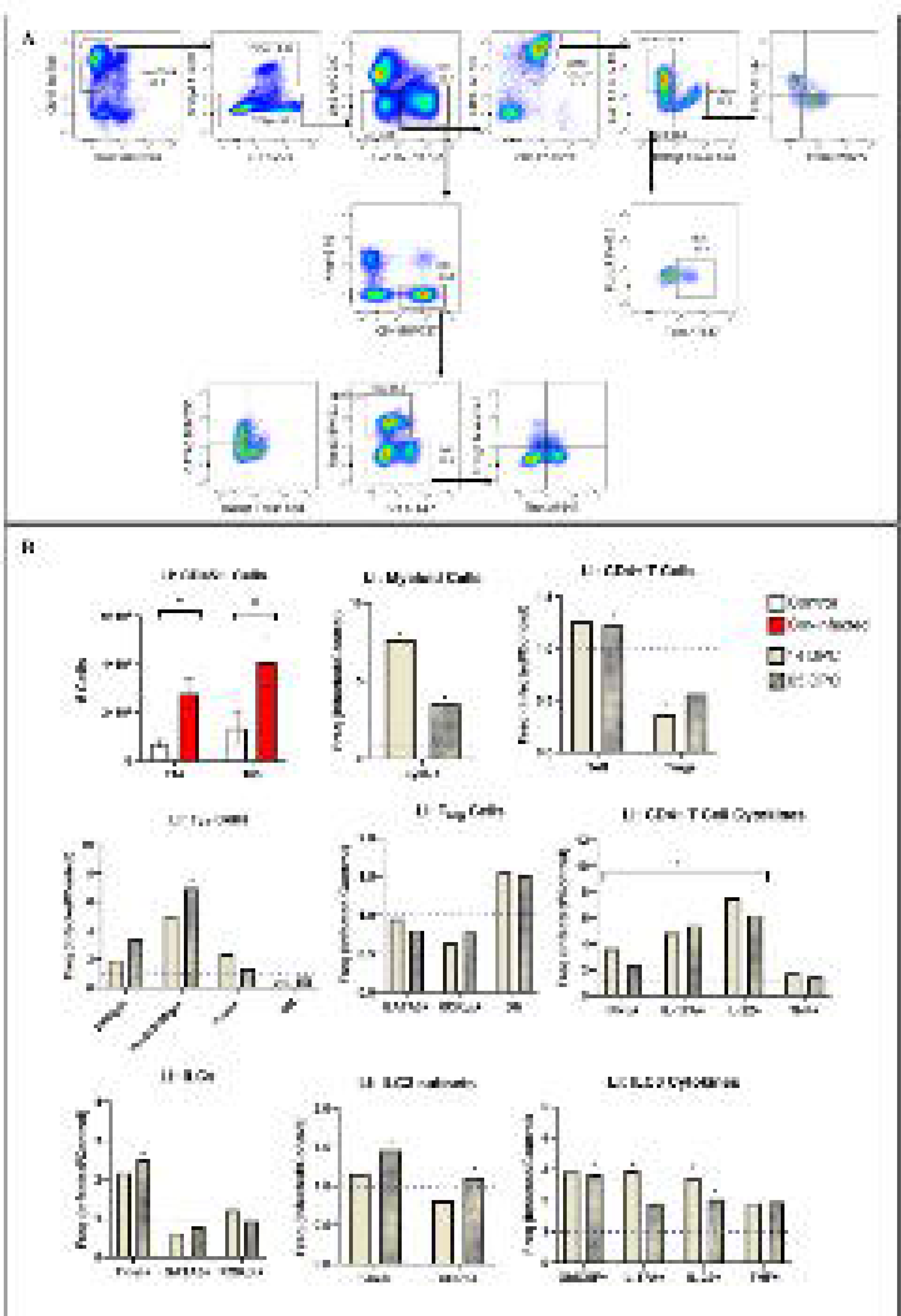
Immunophenotyping of large intestinal immune cells during early and late stages of Cm infection in C57BL/6J (B6) mice. (A) Gating strategy defining the immune subpopulations of the large intestinal cells after excluding debris, doublets, and dead cells. Immune cells were first identified as CD45^+^ cells. T cells were then identified as Lin1^+^, and ILCs as Lin1^-^. CD4 T cells were identified by expression of CD4, and further classified as effector (Tbet) or regulatory (Foxp3) CD4^+^ T cells. ILC3s were identified as CD90 and CD127^+^. Data displayed are from B6 mice at 14 DPC. (B) Representative flow cytometric analyses from CD4 T cell and ILC3 subsets. There were significantly more total immune cells (CD45^+^) in the large intestine of infected mice compared to uninfected controls at both 14 and 63 DPC. Additionally, the cells isolated from the large intestine of infected B6 mice demonstrated increased frequencies of several CD4^+^ T cell and ILC subtypes associated with Th1 and Th17 cell immune responses when compared to uninfected controls. These mice had more frequent T_eff_ CD4^+^ cells (14 and 63 DPC), and fewer T_reg_ cells (14 DPC). These T_eff_ cells more frequently expressed both Tbet and RORγ (63 DPC). Additionally, ILCs and the ILC3 subtype more frequently expressed T-bet (63 DPC), and the ILC3 subtype also more frequently expressed CCR6 (63 DPC). Finally, the large intestine immune cells more frequently expressed several cytokines (with ILC3 having more frequent GMCSF at 63 DPC, IL- 17A at 14 DPC, and IL-22 at both 14 and 63 DPC, and CD4^+^ T cells have more frequent IFNγ, IL17A, and IL22 at both 14 and 63 DPC). Data represent the frequencies of specific cell populations as compared with the parent population and are displayed here as proportions of average values of infected as compared to control mice (n=6 mice/group). Dotted line (1.0) indicates no difference in frequency of detection between control and infected mice. Values above this line represent an increased frequency of detection in infected mice and values below this line represent a decreased frequency of detection in infected mice. macro = macrophage; mono = monocyte; neut = neutrophil; EM = effector memory cell; TIC = total immune cells. *=p≤0.05.

A summary of immunophenotyping results is provided in Table 3.

**Table 3.**
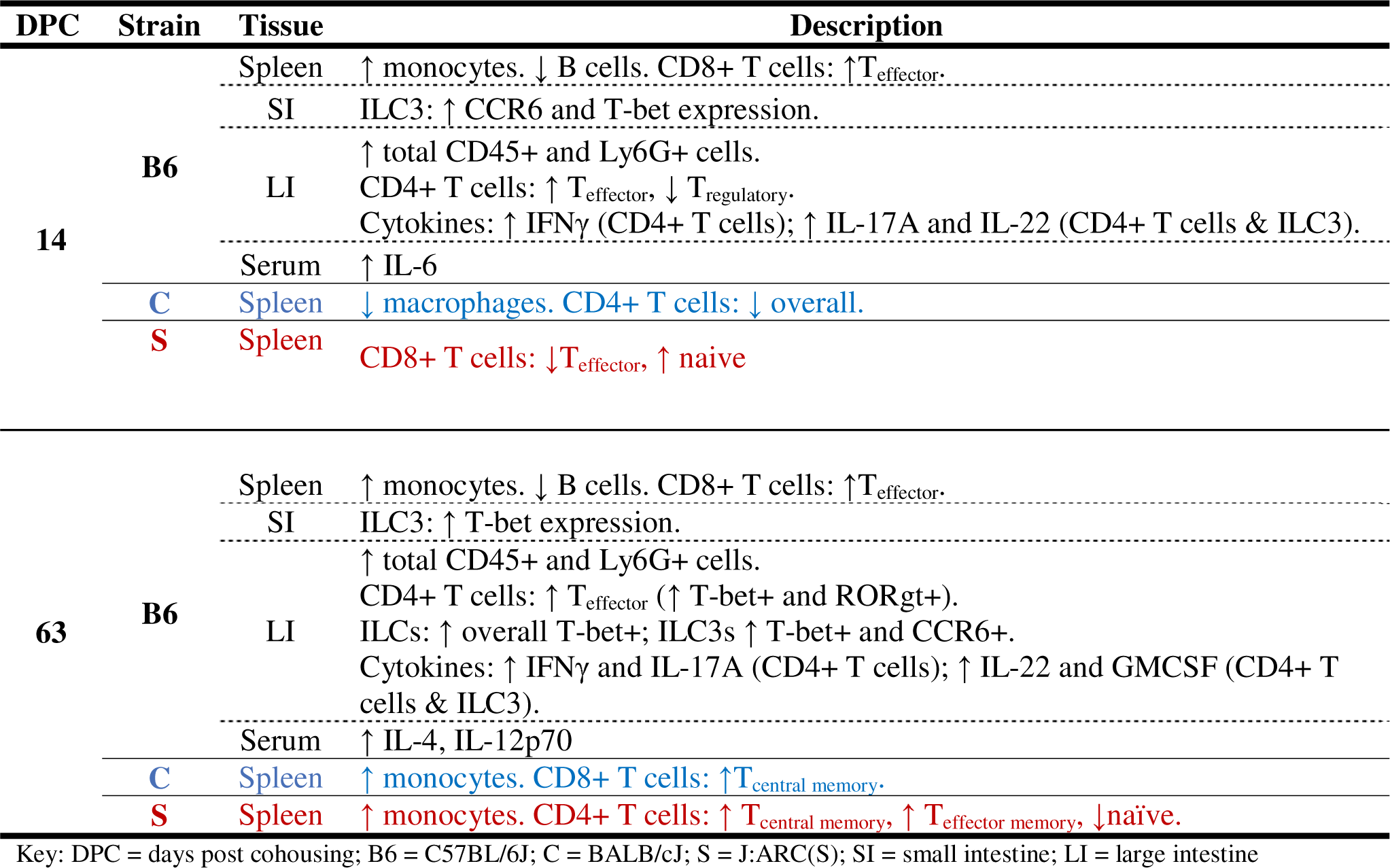
Summary of immunophenotyping results from the spleen (C57BL/6J, BALB/cJ, and J:ARC[S]), small and large intestines, and sera (C57BL/6J) during Cm infection.

### Cytokine secretion assays

Concentrations of the cytokines assayed are provided in Figure 10, and summarized in Table 3. Due to the limited sample size, there were few statistically significant differences noted between Cm-infected and control B6 mice. The increases were consistent with inflammation skewed toward, but not restricted to a Th1 response. At 14 DPC, Cm-infected mice demonstrated an increased expression of the pro-inflammatory cytokine IL-6. At 63 DPC, Cm-infected mice demonstrated increased expression of both Th2 cell-associated IL-4 and Th-1 cell associated IL-12p70 with a more robust increase in the latter. Additional trends observed included increases in IL-2 at 14 DPC, IL-6, IL-10, and MCP-1 at 63 DPC, and GM-CSF, IL-1β, and TNFα at both 14 and 63 DPC. Circulating concentrations of IL-4, IL-10, and IL-12p70 also increased temporally in Cm-infected mice; the concentrations of these cytokines were statistically higher at 63 DPC as compared to 14 DPC. A similar trend was observed in concentration of IL-1β.

**Figure 10.**
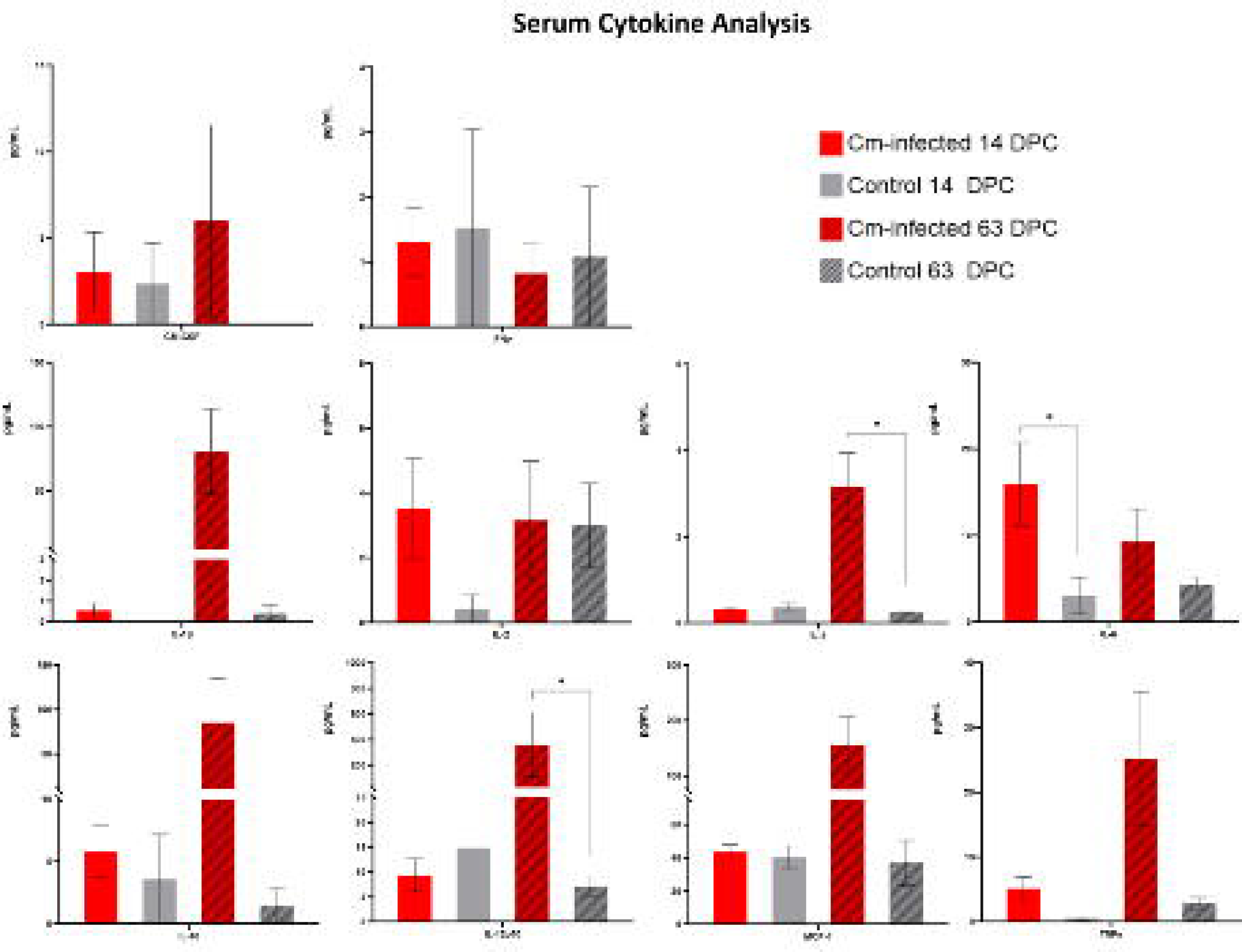
Cytokine expression in Cm-infected C57BL/6J mice at 14 and 63 DPC. Cm-infected mice express significantly more IL-6 than uninfected counterparts at 14 DPC, while at 63 DPC, Cm-infected mice express greater levels of both IL-4 (associated with Th2 responses) and IL-12p70 (associated with Th1 responses) as compared to uninfected controls. IL-12p70 is markedly elevated. Additionally, Cm-infected mice exhibited an increase in IL-4, IL-10, and IL-12p70 from 14 DPC to 63 DPC. Data represents a single experiment with 6 mice/group. Bars represent the SEM. * = p≤0.05.

## Discussion

The present study was conducted to determine the impact of Cm infection on the murine host’s health and biology, as well as to characterize the temporal aspects of infection following infection by the presumptive natural fecal/oral route and the route by which research mice likely become infected. The study’s findings are of importance as it has become clear that Cm is a common component of the mouse’s gastrointestinal microbiota in many research colonies, and commonly utilized immunocompetent inbred and outbred mice develop a chronic, likely life-long, infection of the gastrointestinal tract. Further, while Cm-infected immunocompetent mice do not develop clinical disease, they mount long-lasting local and systemic immune responses.^42,66,69^ The study also demonstrated the potential for mice shedding Cm to efficiently infect other mice through direct contact as well as characterized the tissues where the bacterium localizes and the microscopic changes associated with Cm infection. Collectively, these findings highlight Cm’s potential impact on research mice, especially those utilized in studies investigating gastrointestinal tract physiology and immunology, and provides further evidence that Cm should be included among the agents excluded from research colonies.

Whether inoculated via gavage or cohoused with Cm-shedding mice, all mice in this study were colonized with Cm by 21 DPI/DPC as evidenced by both fecal qPCR positivity and Cm MOMP antigen immunolabeling of gut epithelium. Following cohousing, transmission occurred despite some of the C mice shedding feces with relatively low copy numbers (under 1,000) at the time of cohousing, as compared with peak shedding between 7- and 30-DPI where copy numbers exceeded 100,000 copies. Importantly, chronically infected mice, even with comparatively lower Cm burden, likely act as reservoirs and pose a continuing risk to Cm-free mice within a colony. This result is consistent with recent experimental data suggesting Cm’s ID_50_ is between 100 and 1,000 IFU.^66^ Soiled bedding also serves as an infectious source as evidenced by robust detection of Cm among institutional soiled bedding sentinels.^35,39^ This is in contrast to Cm-contaminated cages, free of Cm-contaminated bedding and feces, which did not lead to transmission of Cm to naïve mice.^39^

Once infected, all B6, C, and S mice shed Cm in feces throughout the duration of the study. We have documented that orogastrically inoculated C mice shed Cm as long as 394 days post-gavage (data not shown). Interestingly, colonization was delayed in the outbred S mice; Cm MOMP antigen immunolabeling was first detected at 21 DPC whereas fecal qPCR positivity was not detected until 35 DPC. While a sample collection error resulted in loss of fecal qPCR data on 21 DPC, the observation that Cm MOMP antigen immunolabeling was always associated with fecal qPCR positivity suggests it is highly likely that S mice did not begin to shed Cm until at least after 14 DPC. Precisely why the outbred mice demonstrated delayed colonization is unclear, however their genetic diversity may result in differences in innate- and adaptive immune responses, epithelial cell receptor differences, dissimilar microbiota composition, and/or delayed consumption of infected Cm fecal pellets from C mice. Recent studies investigating reproducibility in murine infectious disease models have demonstrated that outbred stocks generally mount more robust inflammatory responses altering susceptibility to microbial infection when compared to inbred strains.^3,12,31,38,59^ Outbred mice responded much more robustly to vaccination to *Francisella tularensis* producing more IFNγ and IL-17 than inbred strains.^53^ Another study looking at the immune response before and after bone marrow transplantation described greater baseline levels of CD4^+^ T cells in outbred compared to inbred mice.^21^ Given that these cytokines and T cells are well characterized as crucial for the control of Cm infection, the increased levels observed in outbred mice could have conceivably delayed colonization. While the mechanisms as to how Cm attaches to host cells have not yet been elucidated, there are numerous studies implicating host glycosaminoglycan (GAG) receptors, protein disulfide isomerase (PDI), and other factors affecting cell membrane sulfation in general chlamydial attachment to eukaryotic cells.^17,45,68^ The allelic variability inherent in outbred mice may conceivably yield phenotypic differences in the number of these receptors and/or enzymes present. Despite the temporal difference in colonization observed, all strains/stocks shed similar amounts of nucleic acid at both peak and during steady-state shedding between 63 and 180 DPC. While life-long infection and shedding was not investigated in this study, we believe this is likely based on the current study and findings in experimental models of Cm infection in mice.^66^

Cm was readily detected by fecal qPCR, Cm MOMP immunolabeling, and mRNA ISH. No mice presented with MOMP immunolabeling nor mRNA ISH without concomitant fecal Cm qPCR positivity. However, at early timepoints of infection, mice were qPCR positive but were negative using IHC and ISH. There were 5 Cm PCR positive mice (n=2 B6 and 3 S) that had either absent or inconclusive intestinal tract IHC immunolabeling. In all of these cases, Cm nucleic acid could be detected in gut epithelia using ISH demonstrating that qPCR offers a robust, non-invasive, method of screening for Cm infection. IHC and ISH, on the other hand, offer utility when attempting to evaluate tissue tropism. These tools may be particularly useful when dealing with spontaneous infections in immunodeficient mice presenting with clinical disease.^34,51^ Additionally, they have utility in evaluating colonization of mucosal tissues given the similarity in appearance between goblet cells and IB.

While clinical disease was not appreciated in this study in any infected cohort, there were histologic features associated with Cm infection. With select exceptions (the detection of IB and MOMP immunostaining in the nasopharynx of a B6 mouse and in the stomach of a C mouse), Cm was only detected in the small and large intestine. This is consistent with the presumption that these are the sites of infection and replication in naturally infected mice. Additionally, occasional, minimal-to-mild enterotyphlocolitis was observed, however the significance of this finding is unclear as this inflammation was noted in both Cm-infected and control mice, albeit it was more frequent and of greater magnitude in the former. Inflammatory infiltrates were not consistently found in association with IB and/or Cm MOMP immunostaining and they were observed more frequently at later timepoints. While these mice were not of an advanced age, enteritis and colitis have been reported as background lesions associated with aging secondary to loss of epithelial barrier and/or systemic inflammatory changes.^29,55^ While the observed enterotyphlocolitis may be an age-related incidental change, we cannot absolutely exclude an association with Cm infection.

It is critical to recognize that while clinical disease does not develop in immunocompetent mice, disseminated and sometimes fatal disease can develop in immunocompromised strains.^34,35,42,51,66^ Immunocompetent mice appear able to control Cm infection, preventing dissemination to other tissues and the development of clinically evident disease, however they remain unable to eliminate the bacterium. These findings are consistent with previous oral experimental infections of Cm.^44,62^ This phenotype may be attributable to the formation of non-replicating “aberrant bodies,” downregulation of the immune response/development of host tolerance, and/or bacterial survival mechanisms.^44,69^

Cm was only detected in epithelia and was generally restricted to epithelium overlying GALT, which became hyperplastic most likely a result of chronic immune stimulation. Previous studies investigating mucosal immune responses to enteric bacteria have shown that both dendritic cells and microfold (M) cells sample antigens and bacteria from the intestinal lumen, which ultimately results in both priming and differentiating T and B cell responses in the GALT.^32^ Studies investigating another nonpathogenic intestinal microbe, segmented filamentous bacteria (SFB) describe the induction of both Th17 cells and IL-22 producing CD4^+^ cells in response to colonization.^14,46,49^ SFB exhibits a similar preferential colonization of epithelial cells overlying GALT, attributable to the induction of T cells described above, enhanced production of IgA and subsequent expansion of germinal centers and B cells, and/or production of yet unidentified growth factors.^14,24^ The unique antigen-sampling characteristic of M cells is also utilized by *Salmonella, Yersenia,* and *Shigella* spp.^26^ These pathogenic bacteria have specialized virulence factors and secretion systems to promote mucosal invasion via M cells and induce mucosal immune responses in the gastrointestinal tract.^47^ While the focus of this study was not to determine the cellular tropism of Cm in the intestines, the increased MOMP immunostaining in the surface epithelium overlying the GALT suggests that intestinal epithelial cells may play a role in initiating and/or modulating local and/or systemic immune responses. It was surprising to find that the S mice evaluated at 21 DPC did not have GALT hyperplasia and the feature was occasionally seen in uninfected controls. Regarding the former, it is possible that these S mice did have hyperplastic GALT, however these regions were not captured in the histologic sections examined. It is also plausible that the genetic and microbiota differences in the outbred S stock, which began shedding Cm later than the inbred strains, had delayed and/or led to a disparate immune response to Cm. The GALT hyperplasia detected in uninfected controls was infrequent and less pronounced, and may have resulted from a shift in microbiota with the addition of the cohoused C mice.^56^ While it could be a result of colonization with another infectious agent, this is highly unlikely based on the study design and the husbandry methods employed.^19,63^

The B6, C, and S mice demonstrated significant changes in immunophenotype in the spleen at both the early (14 DPC) and late (63 DPC) timepoints. Monocytes in the spleens of all infected mouse strains are likely enriched in this organ in response to a persistent Cm infection of the host’s gastrointestinal tract. The elevated levels in the monocyte chemoattractant protein-1 (MCP-1) at 63 DPC suggests that monocytes were recruited to the sites of infection. The spleen represents an important reservoir of Ly6C^high^ monocytes, which are typically recruited to sites of infection and inflammation.^54^ Given that monocytes also act as antigen-presenting cells and secrete cytokines in response to *C. trachomatis* infection in vitro, it is possible they prime adaptive immune responses in the T-cell zones of the white splenic pulp in Cm-infected mice.^27^ While Cm inclusions were not noted in the spleen, this organ seems to play a role in the hematogenous spread of Cm from the spleen or extraintestinal tissues to the gastrointestinal trats of mice.^21,70^ Additional immunophenotypic changes in the spleen were noted in T cell subtypes, some of which were significantly different following Cm infection. These changes were consistent with the Th1 cell immune response which Cm is known to induce, and the mouse strains/stock utilized.^7,42^ For example, B6 mice which are known to mount a Th1 skewed response had an increase in “active” CD4+ T cells (via increased effector subtype, T_eff_). Conversely, C mice which are known to mount a Th2 skewed response had fewer CD4+ T cells while outbred S mice presented with increases in both CD4+ and CD8+ responses, consistent with both a Th1 and Th2 response. While the clinical implications of these changes are unclear, researchers evaluating immune responses should consider the impact of Cm infection in their model.

Given that Cm colonizes the gastrointestinal tract and has been demonstrated to primarily elicit a Th1 immune response, we sought to further investigate the large intestinal immune responses of B6 mice following infection with Cm.^62,66^ The local responses were much more robust with both greater numbers of immune cells (CD45^+^), myeloid cells (Ly6G^+^), and effector (activated) CD4^+^ T cells in both early and late timepoints of Cm infection. CD4^+^ T_eff_ were frequently positive for Tbet and RORγ, and produced IFNγ, IL-17A, and IL-22 indicating both a Th1 cell response and Th17 cell activation. These findings demonstrate that both Th1 and Th17 cell responses are induced during Cm infection of the large intestine. In addition, our findings are consistent with recent studies investigating the role of Th1 cell immunity in an experimental model of Cm gastrointestinal infection.^28,64^ However, antigen-specific CD4^+^ T cells were not required for the clearance of Cm in the large intestine of B6 mice, suggesting alternative immune responses may occur in the large intestinal lamina propria in response to Cm infection. We also observed changes in ILC3 responses as evidenced by expression of Tbet and CCR6, which is consistent with recent findings in which innate lymphoid cell responses, especially ILC3, were shown to be a critical component of the immune response to Cm.^20^ ILC3 are considered tissue resident lymphoid cells that are important regulators of intestinal homeostasis between the host and commensal bacteria, and help orchestrate mucosal immune responses to attaching and effacing pathogens, especially bacteria disrupting the intestinal epithelial barrier.^40^

Both the CD4^+^ T cells and ILC3s in the intestinal tissue of B6 mice produced increased pro-inflammatory cytokines when compared to uninfected controls. CD4+ T cells produced increased IFNγ, whereas both ILC3s and CD4+ T cells produced both IL-17 and IL-22 (associated with Th17 responses).^25^ Th17 cells are important in the recruitment of neutrophils to sites of inflammation via IL-17A. While circulating IL-17A was not evaluated in this study, our immunophenotyping data showed a significant increase in Ly6G^+^ granulocytes in the large intestine at 14 and 63 DPC, suggesting a shift in myeloid cells during Cm intestinal inflammation. IL-22 is strongly associated with the maintenance of gut-mucosal immunity and stimulates epithelial cells to produce antimicrobial substances.^33^ Experimentally, Cm has been shown to promote the expression of both IL-17 and IL-22.^15,62^

IL-6 (associated with T-cell survival, downregulation of T_regs_, and as noted is protective against chlamydial infection) and IL-12 (which functions in tandem with IFNγ) were the principal elevated circulating pro-inflammatory cytokines.^65^ We recently reported that mice deficient in IL-12 activity developed fulminant, fatal disease after Cm infection, supporting the important role this cytokine plays in response to infection with Cm.^34^ Additionally, elevations in anti-inflammatory cytokines IL-4 (associated with the differentiation and activity of both B and T cells) and IL-10 (associated with downregulation of the overall host immune response) were observed.^65^ While the role of IL-4, which is classically associated with a Th2 cell response, has not been fully described in chlamydial infections, there is evidence that infection does induce its production in humans.^65^ IL-10 is also important in modulating host response to chlamydial infections in humans.^65^ Interestingly, and as opposed to what was observed in the gastrointestinal tract, increases in circulating IFNγ were not observed. Given the difference in both the magnitude and cellular composition of immune responses observed in spleen vs. gastrointestinal tract, this observation may, in part, reflect a diminished systemic response. However, the relatively small sample size in this study warrants further investigation. The temporal increases in IL-4, IL-10, and IL-12p70 likely reflects the continuing host immune response to Cm infection.

The immunophenotyping we conducted suggests that Cm could have a significant impact on experimental models, especially those involving the gastrointestinal tract. To this point, experimental Cm infection has been shown to negate the ability of DSS to induce colitis through the induction of IL-22.^63^ There are other commonly used mouse models of gastrointestinal disease and/or physiology that may also be impacted if mice were to be colonized with Cm. Infectious disease models, including *Citrobacter rodentium* or *Trichuris muris,* would likely be impacted as Cm alters TLR responses, especially TLR-2 and 3.^5,8^ As TLR-2 protects against *C. rodentium* infection, and is implicated in the host response to *T. muris* infection via mediation of gastrointestinal serotonin production, Cm may confound the experimental model by lessening the pathology following the administration of either.^16,61,67^ Further, as the microbiota and specific infectious agents are known to impact gastrointestinal tumorigenesis by either promoting or suppressing tumor development, Cm’s impact should also be considered.^11,36,71^ Further investigation is warranted to assess the impact of Cm on the aforementioned models. Finally, as immunocompetent mice remain subclinical and infection is persistent, and the infectious dose is relatively low, Cm-infected mice also present a significant risk to immunocompromised mice. Considering the results of this study and the findings in experimental Cm infectivity studies, we posit that testing for Cm should be included among the agents tested for in a colony health monitoring program and Cm should ideally be excluded from research mouse colonies.

## Abbreviations

Cm: *Chlamydia muridarum*
EB: elementary body
RB: reticulate body
IB: inclusion body
C: BALB/cJ
B6: C57Bl/6J
TLR: toll-like receptor
IFU: inclusion forming units
GEM: genetically engineered mouse
NSG: NOD.Cg-*Prkdc*^scid^ Il2rg^tm1Wjl^/SzJ
S: J:ARC(S)
DPC: days post-cohousing
IHC: immunohistochemistry
ISH: in-situ hybridization
MOMP: major outer membrane protein
DPI: days post-infection
BCS: body condition score
GALT: gastrointestinal lymphoid tissue
RORγ: retinoid-related orphan receptor
ILC: innate lymphoid cell
GMCSF: granulocyte macrophage colony stimulating factor
MCP-1: monocyte chemoattractant protein 1
T-bet: T-box transcription factor

## Acknowledgements

MSK Core Facilities are supported by MSK’s NCI Cancer Center Support Grant P30 CA008748.

We thank Maria Jiao and the staff of the Laboratory of Comparative Pathology for their technical assistance with IHC and ISH, Jacqueline Candelier, Simona Bekker, John D’Allara, and Xiaobin Wang for their technical assistance with anatomic and clinical pathology; Morganne Campbell, Abigail Michelson, and Glory Leung for their assistance with data collection; and, Michelle Eckstein with figure creation.

